# Connecting multiway enhancer-promoter interactions to changes in gene expression in cancer

**DOI:** 10.64898/2026.04.25.720760

**Authors:** Kiran Kumari, Sucheol Shin, Guang Shi, Kathleen S. Metz Reed, Tom Misteli, D. Thirumalai

**Author notes:** These authors contributed equally.

## Abstract

Hi-C, and more recently Micro-C, experiments suggest that genome organization is altered as normal cells become cancerous, often resulting in aberrant gene expression. In prostate cancer, there is evidence that large topologically associating domains in normal cells split and are accompanied by shifts in the epigenetic marks from inactive to active states. However, it is unclear whether changes in genome organization alone can account for the gene expression increase in cancer cells. By combining polymer physics concepts and data-driven modeling, we first calculated an ensemble of three-dimensional (3D) chromatin structures using only the 2D contact map as input. The enhancer-promoter (E–P) distance distributions for the overexpressed Androgen Receptor (AR) and Forkhead Box Protein A1 (FOXA1) are broad, with mean values exceeding the threshold for direct E–P contact. Importantly, the average E–P distances decrease in the cancer cells, compared to normal cells in the *AR* locus, whereas in *FOXA1* they are roughly constant or increase modestly. Similarly, the number of multiway contacts increases as cancer progresses across cancer cell lines in the *AR* locus. In contrast, the number of multiway contacts in *FOXA1* is similar in normal and cancer cells. Because the 3D characteristics do not explain the enhanced gene expression in cancer cells, we developed Activity-by-Multiway-Contact (AMC) Model that integrates the multiway contacts with enhancer biochemical activity. The AMC model provides a plausible mechanism for the overexpression of both *AR* and *FOXA1* in prostate cancer. Moreover, using Micro-C data for breast cancer, we show that the number of multiway enhancer-promoter contacts increases in four of five genes studied. When multiway contacts are combined with biochemical activity, changes in gene expression found in the experiment, positively correlate with the AMC score. The predictions of the AMC model not only account for overexpression of genes in prostate and breast cancer but also provide a basis for understanding gene expression variations in other genes as well.

## I. INTRODUCTION

Several studies have used chromosome conformation capture (Hi-C and its variants) experiments to assess the changes in the chromatin organization as normal cells become cancerous, presumably due to the accumulation of somatic mutations [1–8]. The alterations in the genome organization are typically accompanied by aberrant enhancement of the expression levels of a number of genes [9–17]. Until recently [18, 19], the structural changes due to the epigenetic alterations that occur as the normal epithelium becomes cancerous, have been mostly inferred using in situ Hi-C maps, which provide insightful data on the relative contact frequency that two genomically separated loci are in spatial proximity. The Hi-C and more recently Micro-C and related experiments on both normal and cancer cells have established that at the ensemble level (averaging over several million cells), genome organization may be described using two major length scales [20]. On a several megabase (Mb) scale, euchromatin (active or A) and heterochromatin (inactive or B) loci phase separate, which is manifested as checkerboard patterns in the Hi-C contact map [21]. Topologically associating domains (TADs) are prominent on the scale of tens of kb to a few (up to 5Mb) Mb and appear along the diagonal of the contact maps [22]. The TADs, which may be thought of as genome folding units, are important for the following reasons: (1) Pairwise contacts are enriched within TADs. Therefore, the boundaries between TADs likely serve to insulate or restrict inter-TAD interactions. (2) In a number of instances, promoters and the associated enhancers are sequestered within TADs, which could facilitate gene activation. (3) TADs are believed to be conserved across species, although this assertion is not firmly established [23, 24].

The structures underlying the TADs, which are determined by the epigenome profiles, could be used to interpret the differences between cancer and normal cells. In particular, the alterations in the gene regulation in prostate cancer, the focus of the current study, have been analyzed using Hi-C contact maps combined with RNA-seq, DNA FISH and other measures of histone modifications. Let us first summarize the major conclusions of the Hi-C based studies on prostate cancer [4, 5] that are relevant to our computational study. (1) The mean genomic length of TADs is smaller in prostate cancer cell lines than in normal cells. The decrease in the length is typically accompanied by TAD splitting a process in which a large TAD in the normal cell partitions into two or more smaller TADs in the cancer cells. (2) There are several changes in the epigenetic states involving transition from heterochromatin (gene poor or B) to euchromatin (gene rich or A) in multiple genes. Among the large number of genes that are sequestered within the altered TADs a substantial fraction has enhanced expression in the cancer cells. The B to A epigenetic change provides a ready explanation for the enhanced gene expression in cancer cells. (3) Interestingly, TADs that have common boundaries in both the normal and cancer cells are also accompanied by compartment changes. It is notable that genes in the common TADs also show increased gene expression as a consequence of a change in the epigenetic state from inactive (H3K27me3) in normal cells to active (H3K36me3) state in cancer cells. This finding shows the increase in gene expression in the prostate cancer cells is related to the epigenome. However, the relationship between compartment alterations [5], the three-dimensional (3D) chromatin structural changes, and the strength of enhancer activity is unknown. Linking 3D structural changes between normal and prostate cancer cells to gene expression enhancement requires direct imaging data [25] or a computational method [26, 27] that can convert the two-dimensional contact map to 3D coordinates of the chromosomes.

Here, we go beyond 2D contact maps by utilizing the HIPPS method [28], which allows us to generate an ensemble of three-dimensional chromatin structures directly from population-averaged Hi-C data. By converting the contact probabilities into 3D spatial coordinates, we calculate chromatin structural properties, such as spatial distances and multiway interactions, that are inaccessible from contact maps alone. We applied the computational framework to analyze the structural changes that occur in chromatin as the normal prostate cell (RWPE1) becomes cancerous (22Rv1, C42B, and MDAPCa2b). We focus primarily on the two oncogenes, the Androgen Receptor gene (AR) and Forkhead Box Protein A1 gene (FOXA1) [29], whose expression is substantially increased in prostate cancer. Using the generated 3D ensembles, we calculated the changes in the distribution of enhancer-promoter (E–P) spatial distances and multiway contact in which a promoter interacts with multiple (≥ 2) enhancers simultaneously [30]. Although the E–P distances decrease in *AR* in cancer cells, *FOXA1* exhibits minimal distance changes despite significant overexpression, suggesting that pairwise looping models alone are insufficient to explain regulation [31]. Importantly, the probability of formation of multiway contacts is not sufficient to explain the increase in gene expression across the cell lines, implying that genome structure alone is inadequate to differentiate the activities between normal and cancer cells. To explain the experimental data, we developed the Activity-by-Multiway-Contact (AMC) model, inspired by a previous study [32]. The AMC model integrates multiway contact probabilities, derived from 3D structures with epigenetic data, to successfully predict gene expression changes. The same approach also explains the recent findings in breast cancer progression [33].

## II. RESULTS

### 3D chromatin structures from Hi-C data

Is it possible to understand the changes in the gene expression in terms of the variations in the three-dimensional (3D) chromatin architecture between normal epithelial cells (RWPE1) and prostate cancer cells (MDAPCa2b, C42B, and 22Rv1)? To begin to answer this question, we generated ensembles of 3D chromatin conformations using the HIPPS (Hi-C-polymer-physics-structures) method [28]. Specifically, we constructed the 3D structures from population-averaged Hi-C contact maps for both normal and cancer prostate cell lines [4, 5] (see Methods for details). We applied the HIPPS method to different genomic regions containing overexpressed genes in the cancer cells, including the 63–71 Mb region on ChrX, which houses the Androgen Receptor (AR) gene. The simulated contact maps derived from the inferred structures are in agreement with experimental Hi-C data across all three cell lines (Fig. 1a). Importantly, essential features such as compartment identities and TAD boundaries, and TAD splitting in the cancer cells are recapitulated using the inferred structures (Fig. 1), even though the input in these calculations are only the Hi-C contact maps. Besides the excellent agreement between HIPPS-generated and the experimental contact maps in all the cell lines (Fig. 1a), the computations also correctly account for the splitting of the TADs in C42B and 22Rv1 (Fig. 1b) found in experiments [4]. Although not noted in the Hi-C experiments on MDAPCa2b cells [5, 34], our findings show that TAD splitting occurs in ChrX in the region containing the *AR* locus (see the last panel in Fig. 1b). Finally, the differences in the average distance maps with respect to RWPE1 for the three cancer cells C42B, 22Rv1, and MDAPCa2b in Fig. 1c show that there are substantial variations. However, the distance differences of the enhancers relative to the *AR* gene is only about ≈ 50 nm, which is less than the resolution of the Hi-C experiments [4] (see SI for an estimate of the spatial scale resolution for Hi-C experiment).

**FIG. 1:**
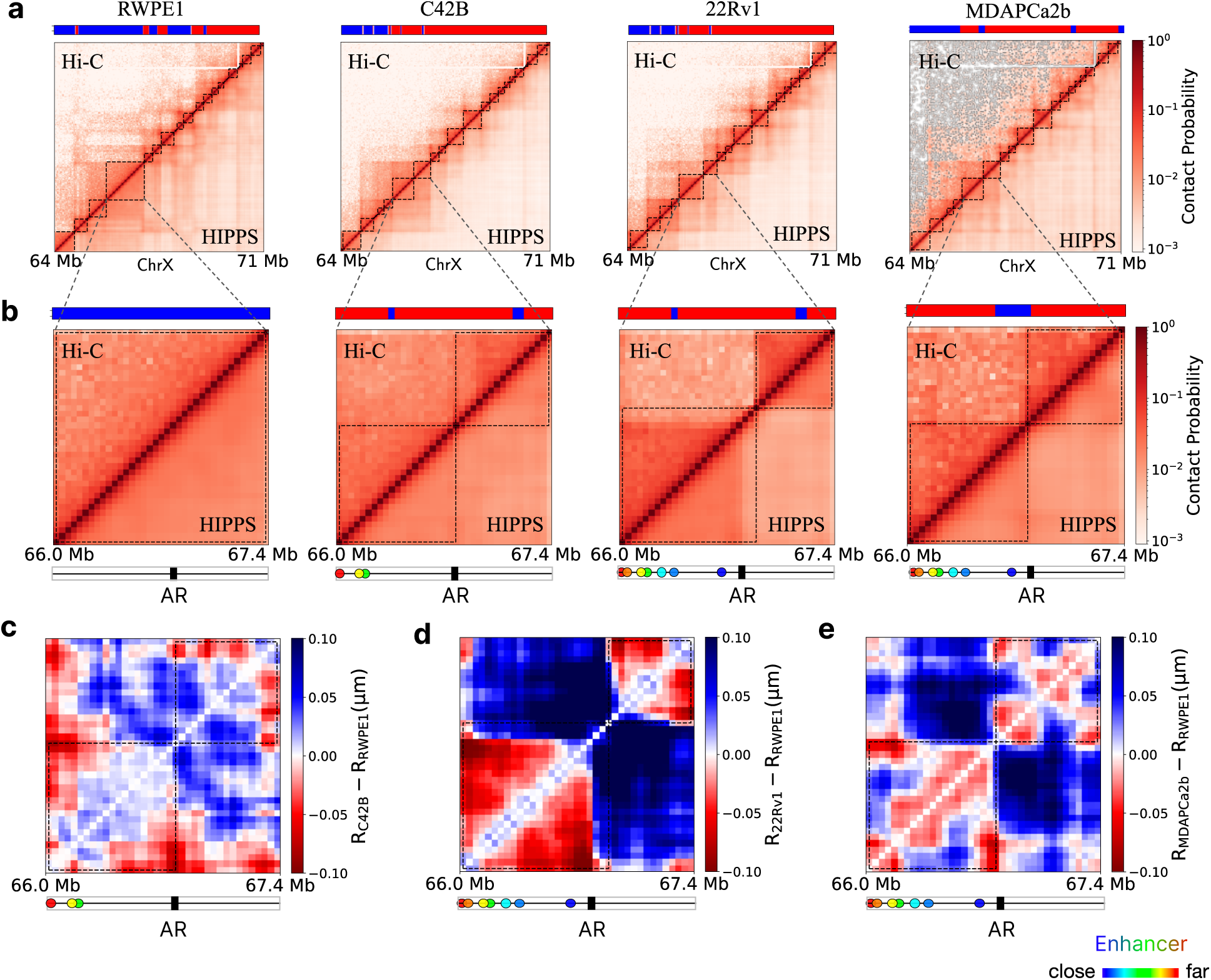
Comparison of calculated and experimental 3D chromatin contacts reveals AR-locus reorganization and TAD splitting in prostate cancer cell lines. (a) Contact maps for the genomic region spanning 64–71 Mb on chromosome X, comparing Hi-C measurements with contact probabilities computed from the 3D structure ensemble reconstructed using HIPPS [28]. Cell lines are indicated above each map. For each cell line, the upper (lower) triangle shows the Hi-C (HIPPS) contact map. Dashed squares along the diagonal indicate TAD boundaries. Compartment assignments inferred from principal component analysis (PCA) of the Hi-C data (A compartment in red, B compartment in blue; see SI Methods) are shown above. (b) Zoom-in to 66.0–67.4 Mb. This region forms a single TAD in RWPE1 but splits into two sub-TADs in the cancer cell lines. The *AR* promoter and associated enhancers are marked below the maps: the promoter is shown as a black rectangle and enhancers as color circles. The blue (red) circle denotes the enhancer closest to (farthest from) the *AR* promoter in genomic distance. (c–e) Differences in the mean distance maps between each cancer cell line and RWPE1: (c) C42B–RWPE1, (d) 22Rv1–RWPE1, and (e) MDAPCa2b–RWPE1.

### Androgen Receptor (AR) gene

To investigate the relationship between changes in the gene expression and the associated alterations in the 3D chromatin structure, as the normal cells become cancerous, we focused on two key genes—Androgen Receptor (AR) located on chromosome X, and Forkhead Box Protein A1 (FOXA1) on chromosome 14. Both these genes play important roles in various stages of prostate cancer. We focused on them because their expression is increased by more than 500-fold in 22Rv1 cancer cells compared to normal RWPE1 cells [4]. Interestingly, although both genes are overexpressed in cancer to the same extent, their three-dimensional chromatin organizations are markedly different, as we show below.

The 1.4 Mb region spanning 66 Mb to 67.4 Mb on chromosome X contains the *AR* gene (Fig. 1b). Hi-C experiments [4, 5, 34] have demonstrated that compartment identity changes are common in prostate cancer cells. In the normal RWPE1 cells, this region is entirely within the inactive compartment, whereas in the cancer cell lines it shifts predominantly to the A compartment, accompanied by TAD splitting (Fig. 1b). Along with compartment changes, TAD splitting also results in the insulation of the *AR* gene from its upstream region, corresponding to the upper TAD in the three cancer cell lines as observed in Fig. 1b. As a result of TAD splitting, it is possible that the insulation mechanism that protects the *AR* locus in the RWPE1 cells may no longer be effective in the cancer cells. This provides a qualitative, but not quantitative 3D structural, explanation for the increase in the gene expression in cancer cells.

The generation of the 3D structures allows us to calculate, among other things, the distance maps (DMs), which give the mean distances between any two loci. The differences in the DMs between the normal and three cancer cells (Fig. 1c) reveal that there are cancer-cell-line-dependent variations in the genome organization. The differences in the DMs between the three cancer cells and RWPE1 are substantial. Fig. 1c shows that the distances between the promoter and the enhancers in the 22Rv1 cell line are closer (distances decrease by nearly 100 nm). In contrast, the decrease in the distances is far less in the C42B cell line. These results show that enhancer–promoter pairs are, on an average, closer in cancer cells than in normal cell. The ensemble of structures determined using the HIPPS method shows *AR* locus exhibits extensive cell line–dependent heterogeneity, reflected by changes of distributions of distances across the distinct cell lines (see SI for details).

### 3D E–P distances in the AR locus decrease in cancer cell line

Having demon strated that there are qualitative, population-averaged changes in the enhancer–promoter distances between normal and cancer cells (Fig. 1c), we next examined the enhancer-specific changes quantitatively. Enhancers, which are thought to be evolutionarily conserved non-coding elements [35], regulate gene expression through complex but poorly understood mechanisms [36, 37]. One is the loop model which posits that enhancers that could be separated from the promoter by large genomic distances (GDs) can be brought to small spatial distances to enable E–P interactions. In the process, the intervening chromatin forms a loop, whose formation is often mediated by architectural protein (CTCF) and the loop-extruding motor cohesin [38–40], bringing transcription factors and co-activators bound to enhancers near the gene promoter, facilitating gene activation. If the looping model holds, then it follows that the E–P distances should decrease in cancer cells relative to the normal cells because the gene expression in the former is enhanced.

To test the looping model, we first calculated the average spatial distances (SDs) between the *AR* promoter and loci *i* (Fig. 2a), measured with respect to the promoter location. Interestingly, the SDs increase but not monotonically, as predicted by standard polymer models often used to rationalize chromatin organization, as the GD increases. In all the cell lines, there is a plateau in ⟨*R_P,i_*⟩ in the neighborhood of the eight enhancers (defined in [4] and listed in Table SI-2) that are upstream of the promoter, which suggests that there could exist favorable E–P interactions. Similar conclusions can be reached from the calculated distances of the loci that are downstream of the promoter (Fig. 2a). A second notable feature is observed downstream of the *AR* promoter, where no enhancers are annotated. In 22Rv1, a locus located ∼ 0.2 Mb from the promoter is farther in 3D (∼ 0.54 *µ*m) than in RWPE1 (∼ 0.40 *µ*m) (Fig. 2a), implying a loss of proximity in the enhancer-poor regions in cancer. This increased separation is consistent with substantial reorganization of chromatin in prostate cancer.

**FIG. 2:**
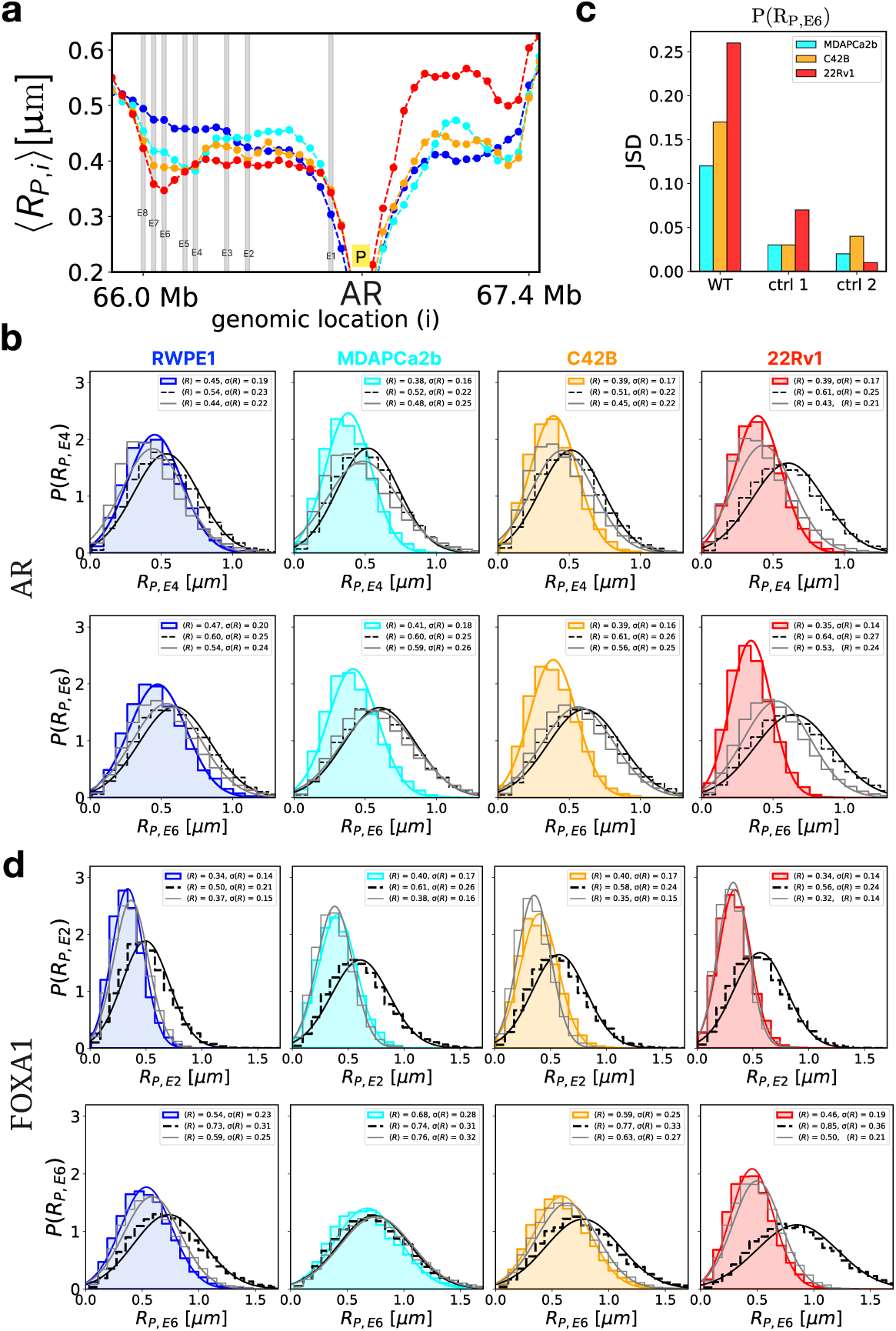
E–P distances for AR and FOXA1 across normal and prostate cancer cell lines. (a) Average distance from the *AR* promoter to other locus in chromosome X region spanning from 66 Mb to 67.4 Mb for the various prostate cell lines (blue: RWPE1, cyan: MDAPCa2b, orange: C42B, red: 22Rv1). Gray vertical lines mark the locations of eight enhancers (E1 to E8). (b) Top panel: Probability distribution of the 3D distance between the *AR* promoter (P) and enhancer E4 across the cell lines. Control 1 (black dashed) shows the promoter–locus distance distribution for a locus at the same genomic distance (GD) from P as E4. Control 2 (gray solid) is the distance distribution for randomly chosen locus pairs within the same chromosomal region whose GD matches the P–E4 GD; because many such pairs exist, we plot the average over all eligible pairs. The bottom panel repeats the top-panel analysis for P–E6 distances, replacing E4 with enhancer E6. (c) Jensen–Shannon divergence (JSD) between the RWPE1 distance distribution *P* (*R_P,E_*_6_) (WT) and the corresponding distributions in the cancer cell lines (C42B, 22Rv1, and MDAPCa2b). For comparison, JSD values for the two controls are also shown. (d) Probability distributions of promoter–enhancer distances for the *FOXA1* gene, shown for two enhancers: E2 and E6. The analysis is identical to the *AR* promoter–enhancer panels, but applied to *FOXA1*. Enhancer locations are provided in SI Fig. 11.

We also compared the E–P distance distributions *P* (*R_P,Ei_*) (*i* = 1*, . . .,* 8) across cell lines, where *R_P,Ei_* is the 3D distance between the *AR* promoter *P* and enhancer *E_i_*. The list of enhancers is in Table SI-2. Unless otherwise noted, the genomic locations of the enhancers (E1–E8) are taken from Rhie *et al.* [4], based on histone modifications in 22Rv1. Because enhancer annotations are cell-type specific, we fixed the set of eight genomic locations in each cell line, including RWPE1. This enables comparisons across cell lines on equal footing (SI Figs. S6 and S7). Across the prostate cancer cell lines, several upstream loci show a clear reduction in the E–P distances relative to RWPE1, most prominently for E4 and E6 (Fig. 2b). The distance distributions are broad: for many loci, the coefficient of variation *σ*(*R*)*/*⟨*R*⟩ ̰> 0.4, indicating substantial cell-to-cell variability. The decrease in mean distance is most pronounced in 22Rv1 (last panels of Fig. 2b). In contrast, for loci beyond E8 (the largest genomic distance from the promoter among E1–E8), promoter–locus distances are similar across all cell lines, demonstrating that the reorganization is localized close to the enhancers rather than genome-wide.

To assess whether the short E–P distances in cancer cells due to mutational effects simply reflect polymer scaling with GD, we constructed two controls. In Control 1, we computed *P* (*R_P,λi_*), where *λ_i_* is a non-enhancer locus chosen such that GD(*P, λ_i_*) = GD(*P, E_i_*). In Control 2, we selected a pair of loci, neither of which is *P* or *E_i_*, whose GD matches that of (*P, E_i_*). For example, in the upper (lower) subpanel of Fig. 2b, Control 1 corresponds to the pair (*P, λ*_4_) [(*P, λ*_6_)], which matches the GD of (*P, E*_4_) [(*P, E*_6_)]. In both controls, the distribution peaks occur at larger 3D distances than for the corresponding E–P pairs. For instance, in 22Rv1 the peak of Control 2 is ∼ 180 nm larger than the peak of *R_P,E_*_6_ . More-over, across all cell lines, the control distributions are consistently broader than *P* (*R_P,E_*_4_) and *P* (*R_P,E_*_6_), supporting the interpretation that enhancer loci in the *AR* gene exhibit unusually frequent close promoter contact beyond what is expected from genomic distance alone.

Fig. 2c shows the Jensen-Shannon Divergence (JSD) between the cancer cell lines and the normal RWPE1 cell line. For *P* (*R_P,E_*_6_), the JSD for MDAPCa2b is relatively low (0.1) and increases to 0.15 for C42B, reaching 0.25 for 22Rv1. In contrast, the JSD values for the control samples remain small, indicating high similarity between the normal and cancer cell lines in these two control cases. Taken together, Fig. 2 suggests a gain in enhancer-promoter proximity within the *AR* locus compared to the controls. Such a conclusion supports the looping model but does not explain the findings in *FOXA1* gene, as we show next.

### Changes in E–P distances at *FOXA1* Gene Locus

We next examined another key gene involved in prostate cancer progression. Mutations in the oncogene *FOXA1* are common in prostate cancer and play an important role in gene regulation. Many overexpressed genes in the cancer cells have the *FOXA1* motif in the promoter regions, indicative of its role in activating gene expression. *FOXA1*, located on chromosome 14 between 36 Mb and 44 Mb, is significantly overexpressed in prostate cancer cell lines, showing a fold increase of ≈ 500 in 22Rv1 and in C42B, compared to normal cell lines [4].

To assess whether the decrease in the average E–P distance observed in the *AR* locus is also present at the *FOXA1* locus, we performed the same analysis. *FOXA1* gene has 8 enhancers (listed in SI Table SI-3). Unlike the *AR* gene, the super-enhancers of *FOXA1* are positioned on both sides of the promoter, extending upstream and downstream. SI Figures 9a and 9b illustrate the Hi-C contact map along with the locations of the enhancers and promoters of *FOXA1*. In addition, SI Figure 9c shows the difference in the average distance map derived from structures generated using HIPPS.

Surprisingly, the analysis of *FOXA1* reveals an entirely different pattern from the *AR* locus. While the *AR* locus shows consistent decreases in E–P distances in cancer cell lines, the *FOXA1* gene locus exhibits only modest changes in pairwise E–P distances despite the dramatic overexpression (500-fold increase) in the cancer cells. The mean ⟨*R_P,E_*_2_⟩ increases by ≈ 60 nm in C42B relative to RWPE1 (Fig. 2d). The distance distributions, *P* (*R_P,E_*_2_) and *P* (*R_P,E_*_6_), associated with enhancers E2 and E6 are broad with substantial dispersions (Fig. 2). Note that enhancer E2 is located upstream of the *FOXA1* promoter, while enhancer E6 is positioned downstream. The distance distributions of all enhancers relative to the promoter, presented in SI Figures 10 and 11, show that the sample range of distances with substantial dispersion. Importantly, the distances are almost the same in all the cell types although there is modest increase involving a few enhancers.

The findings so far point to a few conclusions. For the *AR* locus, E–P distances are shorter than in the control cases for loci pairs at the same genomic distances, even when the gene is transcriptionally inactive, as observed in the *AR* gene in RWPE1. A similar pattern is observed at the *FOXA1* locus. These results are consistent with recent single-cell imaging studies reporting that promoter–enhancer pairs tend to be in closer proximity than non-enhancer pairs separated by comparable genomic distances [31]. In the cancer cell lines, the *AR* locus shows further reduction in E–P distances and a smaller spread in the distance distributions. However, this pattern is not always observed: the *FOXA1* locus exhibits little change in the E–P distances despite the observed 500-fold overexpression, suggesting that alternative mechanisms are operative. Using the calculated 3D coordinates we explored the possibility that multiway enhancer–promoter contacts (defined as multiple enhancers in contact with promoter) might play a dominant role in gene regulation.

The difference between *AR* and *FOXA1* loci suggests that the mechanism of gene expression enhancement in cancer cells is not solely due to pairwise E–P distance changes, which is inconsistent with the loop model. The observation that *FOXA1* shows minimal E–P distance changes despite substantial overexpression while AR shows significant distance changes with similar overexpression (500-fold) indicates that alternative mechanisms, such as multiway enhancer-promoter contacts, might play a more prominent role in certain gene loci. This possibility prompted us to investigate multiway contact patterns rather than relying only on pairwise E–P distance analysis. We also analyzed other genes including *RBM11* (which has a single enhancer) and *SHH* to further explore the diversity of regulatory mechanisms (see SI for more details).

### Multiway enhancer-promoter contacts for *AR* and *FOXA1*

So far, we focused on the pairwise 3D distances between specific pairs of enhancers and promoters. However, a single promoter frequently interacts with multiple enhancers simultaneously, which raises important questions: Is there a particular enhancer or a small number of them that play a crucial role? Must a promoter interact with many enhancers to form hub-like structures to activate a gene? Exploring these questions, which require knowledge of 3D structures at the single-cell level, could provide valuable insights into the mechanisms of gene regulation in normal and cancer cells.

To address these questions, we extended our analysis to determine multiway contacts between promoters and enhancers for both the *AR* and *FOXA1* loci. Specifically, we examined cases in which various enhancers (*n* = 1, 2, *…*8) are in contact with the promoter simultaneously. The contact threshold, *r_c_*, is set at 1.5*σ* = 1.5 × 160 nm= 240 nm, where *σ* represents the monomer size. Using this threshold, the Pearson correlation between the experimental Hi-C data and the contact probability map generated by HIPPS is approximately 0.95 (see SI for details).

*AR locus:* Fig. 3a presents the probability distribution of *n*-way contacts, *P_n_*, between the *AR* promoter and the associated *n* enhancers. The distribution is normalized by the probability that the promoter is not in contact with any enhancer (*P*_0_). There is a clear trend: as *n* increases, *P_n_* decreases but depends on the cell line. This finding is not unexpected because the likelihood of multiple enhancers interacting with the promoter simultaneously should diminish as the number of enhancers increases, principally due to excluded volume effects. Such a behavior is also consistent with general polymer statistics because the probability of forming higher-order (multiway) contacts is expected to decrease rapidly with *n*. Interestingly, the decrease in *P_n_* is steeper in normal cells than in cancer cell lines. For instance, when *n* = 3, the *AR* promoter in cancer cells exhibits a higher probability of interacting with three enhancers compared to normal cells. Among the three cancer cell lines examined, 22Rv1 consistently shows the highest *P_n_* across all values of *n*, coinciding markedly with elevated *AR* expression level.

**FIG. 3:**
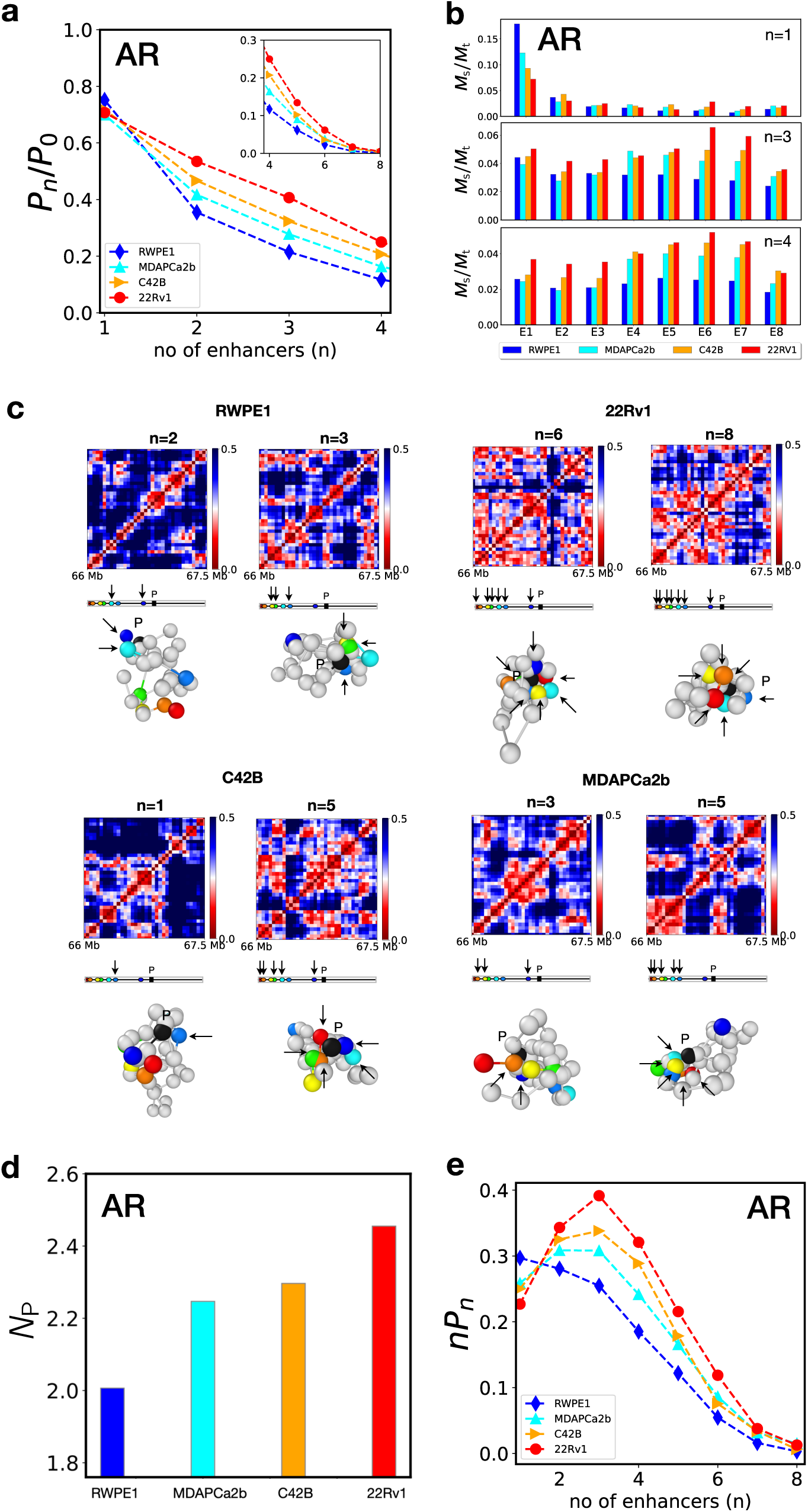
Multiway E–P contacts in *AR* across cell lines. (a) The relative probability, *P_n_/P*_0_, of *n*-way contacts of the promoter with enhancers where *n* is the number of enhancers involved in the contact. The probability that no enhancer is in contact with the promoter is *P*_0_. The values of *P*_0_ are approximately 0.40, 0.37, 0.35, and 0.32 in RWPE1, MDAPCa2b, C42B, 22Rv1, respectively. *P_n_* (*n* = 1, 2, 3*, ..*) is the probability that *n* enhancers are in contact with the promoter. The inset shows *P_n_* for *n* from 4 to 8. (b) Upper panel: The contribution of each enhancer when only one of the eight enhancers forms a binary contact (*n* = 1) with the promoter. *M_s_* is the number of conformations involved in the contact and *M_t_* is the total number of conformations. Middle panel: The same analysis as in the upper panel, but for cases of *n* = 3. Lower panel: The same analysis as in the upper panel, but for cases of *n* = 4. (c) Distance maps for the RWPE1 and three cancer cell lines for *n* number of enhancer contact. Below each distance map, the corresponding structure is displayed. The bead in black is the promoter. The arrows mark the enhancers that are in contact with the promoter. (d) Average number of E–P multiway contacts (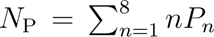) for the *AR* locus in normal and cancer cell lines. (e) Contribution of each *n*-way contact to the total multiway contacts for the *AR* locus across the four cell lines.

We next examined whether different enhancers contribute equally to multiway E–P interactions. We analyzed two representative cases: (i) conformations in which the *AR* promoter interacts with only one of the eight enhancers (*P*_1_), and (ii) conformations in which it interacts with four enhancers simultaneously (*P*_4_). For each enhancer, we calculated the fraction of conformations with a fixed number of E–P contacts (*n* = 1 or *n* = 4), defined as *M_s_/M_t_*, where *M_s_* is the number of conformations involving the enhancer and *M_t_* is the total number of conformations with *n*^th^-order E–P contacts. Fig. 3b clearly shows that, in *P*_1_, enhancer E1 (the genomically closest enhancer to the promoter) is dominant, exhibiting the highest contribution to the multiway contacts across all the cell lines. Notably, in the normal epithelium (RWPE1), the contribution of E1 is the greatest but decreases modestly in the cancer cell lines. For the other enhancers, their contributions are relatively consistent between normal and cancer cell lines. Strikingly, for *n* = 4 (chosen for illustrative purposes), enhancers E4 to E7 become prominent. Two key conclusions emerge: (i) all enhancers make higher contributions in cancer cell lines compared to the normal epithelium, and (ii) enhancers E4 to E7 contribute significantly more than the other others in cancer cell lines.

We then analyzed the average number of enhancers that simultaneously contact the promoter, denoted as *N*_P_, defined as, 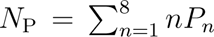 where *n* is the number of enhancers in contact with the promoter and *P_n_* is the corresponding probability. Fig. 3d shows the average number of enhancers simultaneously contacting the promoter (*N*_P_) for each cell line. Among the cell lines, the normal epithelial line RWPE1 has the lowest level of multiway promoter–enhancer contact, with *N*_P_ ≈ 2.0. The cancer cell lines MDAPCa2b and C42B show intermediate levels, with *N*_P_ values between 2.2 and 2.3. Notably, the cancer cell line 22Rv1 has the highest degree of multiway interaction, with *N*_P_ = 2.4, reflecting a particularly dense network of enhancer–promoter contacts that may underlie the elevated *AR* gene expression.

We further examined how contacts involving different numbers of enhancers contribute to the overall average, *N*_P_. The distribution of *nP_n_* for different values of *n* for the normal RWPE1 cell line has a peak at *n* = 1 (Fig. 3e), indicating that higher-order (multiway) contacts are relatively rare. In contrast, in cancer cell lines, the peak shifts to *n* = 3 or *n* = 4, with the most pronounced effect observed in 22Rv1. This pattern suggests the formation of transcriptional hubs in which the *AR* promoter simultaneously engages with up to four enhancers—approximately five-way interactions—that may facilitate its strong transcriptional activity.

*FOXA1:* We then performed a similar structural analysis for the *FOXA1* gene. Fig. 4a shows the *P_n_/P*_0_ ratio for *FOXA1*, which reveals that normal cell RWPE1 and cancer cell 22Rv1 show the most prominent multiway contacts, whereas C42B and MDAPCa2b show similar but significantly less multiway contacts. This is in stark contrast to the *AR* locus, which shows a consistent trend of increasing multiway contacts from RWPE1 to C42B to 22Rv1. Fig. 4b shows the *M_s_/M_t_* ratio for *FOXA1*, demonstrating that RWPE1 and 22Rv1 have more multiway contacts compared to the other cell lines. Fig. 4d shows *N*_P_ across the four cell lines, again confirming that RWPE1 and 22Rv1 have the most prominent multiway enhancer-promoter contacts with higher *N*_P_ values, compared to MDAPCa2b and C42B, consistent with Figs. 4a and b. Finally, Fig. 4e shows *nP_n_* as a function of the number of enhancers (*n*) for *FOXA1*, which also shows the same pattern: RWPE1 (normal) and 22Rv1 (cancer) show similar multiway contacts, more prominent than the other cell lines. These results show that no clear conclusion regarding the role of multiway contacts can be drawn for the *FOXA1* gene.

**FIG. 4:**
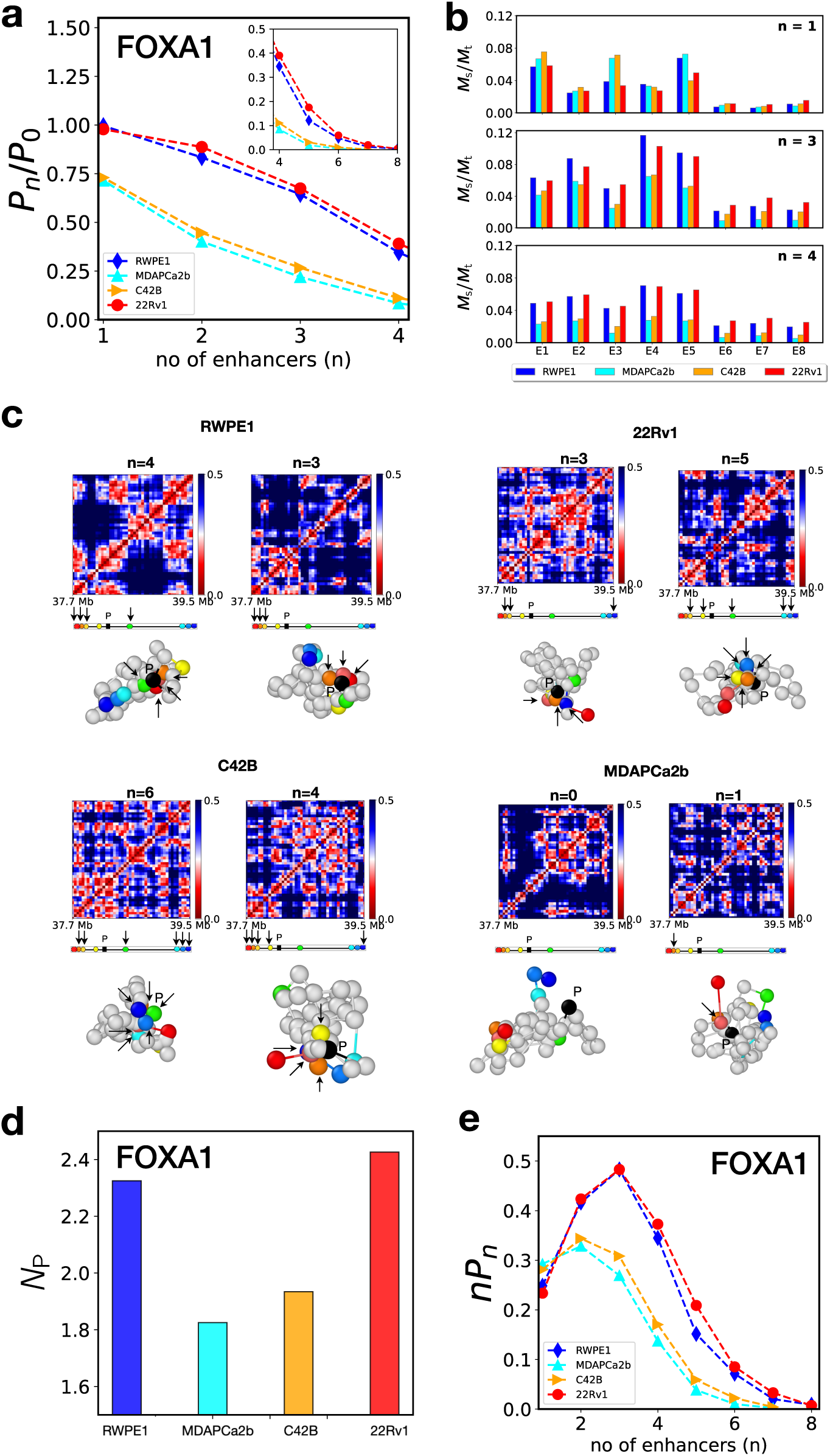
Multiway E–P contacts in *FOXA1* across different cell lines. (a) Relative probability distribution, *P_n_/P*_0_, of *n*-way contacts for the *FOXA1* locus. The values of *P*_0_ are approximately 0.25, 0.41, 0.39, and 0.24 in RWPE1, MDAPCa2b, C42B, 22Rv1, respectively. (b) *M_s_/M_t_* ratio for *FOXA1*, showing the fraction of conformations with multiway contacts. (c) Distance maps for the RWPE1 and three cancer cell lines for *n* number of enhancer contact. Below each distance map, the corresponding structure is displayed. The bead in black is the promoter. The arrows mark the enhancers that are in contact with the promoter. (d) *N*_P_ for *FOXA1* across the four cell lines. (e) Distribution of *nP_n_* as a function of the number of enhancers (*n*) for *FOXA1*, showing the contribution of different multiway contact orders.

The contrasting behavior of the *AR* and *FOXA1* loci suggest that differences in gene expression may not be explained solely by the number or strength of multiway contacts. Although these contacts reflect the structural organization of chromatin, transcriptional regulation also depends on epigenetic factors such as histone modifications. We therefore developed a method that integrates multiway contact data with epigenetic information to investigate how 3D chromatin architecture and epigenetic modification together shape gene expression.

### Activity-by-multiway-contact (AMC) model

Enhancer activity and spatial organization vary across cell types [4]. Fulco *et al.* [32] introduced the Activity-by-Contact (ABC) model to describe enhancer contributions to gene expression. In the ABC model, enhancer activity is measured using an integrated metric that combines epigenetic information (such as DNase-seq, H3K27ac, ChIP-seq) with the E–P contact probabilities derived from Hi-C data. The ABC score of a given enhancer correlates with the change in gene expression upon perturbation of that enhancer [32]. Building on this concept, we introduce the Activity-by-Multiway-Contact (AMC) model, a structural ensemble-based method, to examine the effect of the differences in 3D chromatin architecture on the differential gene expression between normal and cancerous prostate cells.

Unlike the ABC score, which is defined for individual enhancers, the AMC score is defined for a given gene based on the 3D chromatin structure surrounding the locus. For a given 3D chromatin conformation, *ξ* = {***r****^i^*}, the AMC score for gene *g* is given by,

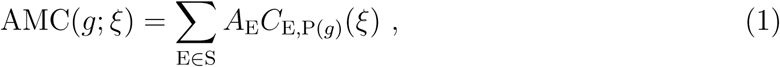

where *A*_E_ is the biochemical activity (specified by epigenetics) of an enhancer, E. P(*g*) is the promoter of gene *g*. S is the set of enhancer candidates. We used H3K27ac ChIP-seq read counts as a measurement of *A*_E_. *C_E,P_*(_*g*_)(*ξ*) is an indicator function that specifies whether enhancer E is in contact with the promoter of gene *g* in conformation *ξ*. It is unity if E is in contact with the promoter (*C_E,P_*(*_g_*)(*ξ*) = 1), and is zero otherwise (*C_E,P_*(*_g_*)(*ξ*) = 0). Note that, if the enhancer activity is a constant, then AMC(*g*; *ξ*) = *n*_P(_*_g_*)(*ξ*) where *n*_P(_*_g_*)(*ξ*) is the number of enhancer candidates in contact with P(*g*), as reported in Figs. 3 and 4. The ensemble average ⟨AMC(*g*)⟩ over the conformational ensemble is equivalent to the ABC score (SI).

We already established that structural information alone is insufficient to understand gene expression differences between normal and cancer cells. The AMC model is implemented by first identifying the most distal enhancers associated with each gene [4]. The genomic region spanning between the furthest upstream and downstream enhancers was then defined as the analysis window, and all loci within this window were included in the summation in Eq. 1. Accordingly, we treated all genomic loci within this window as potential enhancers contributing to the AMC score. Fig. 5a schematically illustrates the enhancer range defined for *AR* and *FOXA1*. Note that if no enhancer is identified for a given gene, then AMC score is not defined. This approach ensures comprehensive coverage of potential regulatory elements while maintaining consistency with the established E–P interaction patterns.

**FIG. 5:**
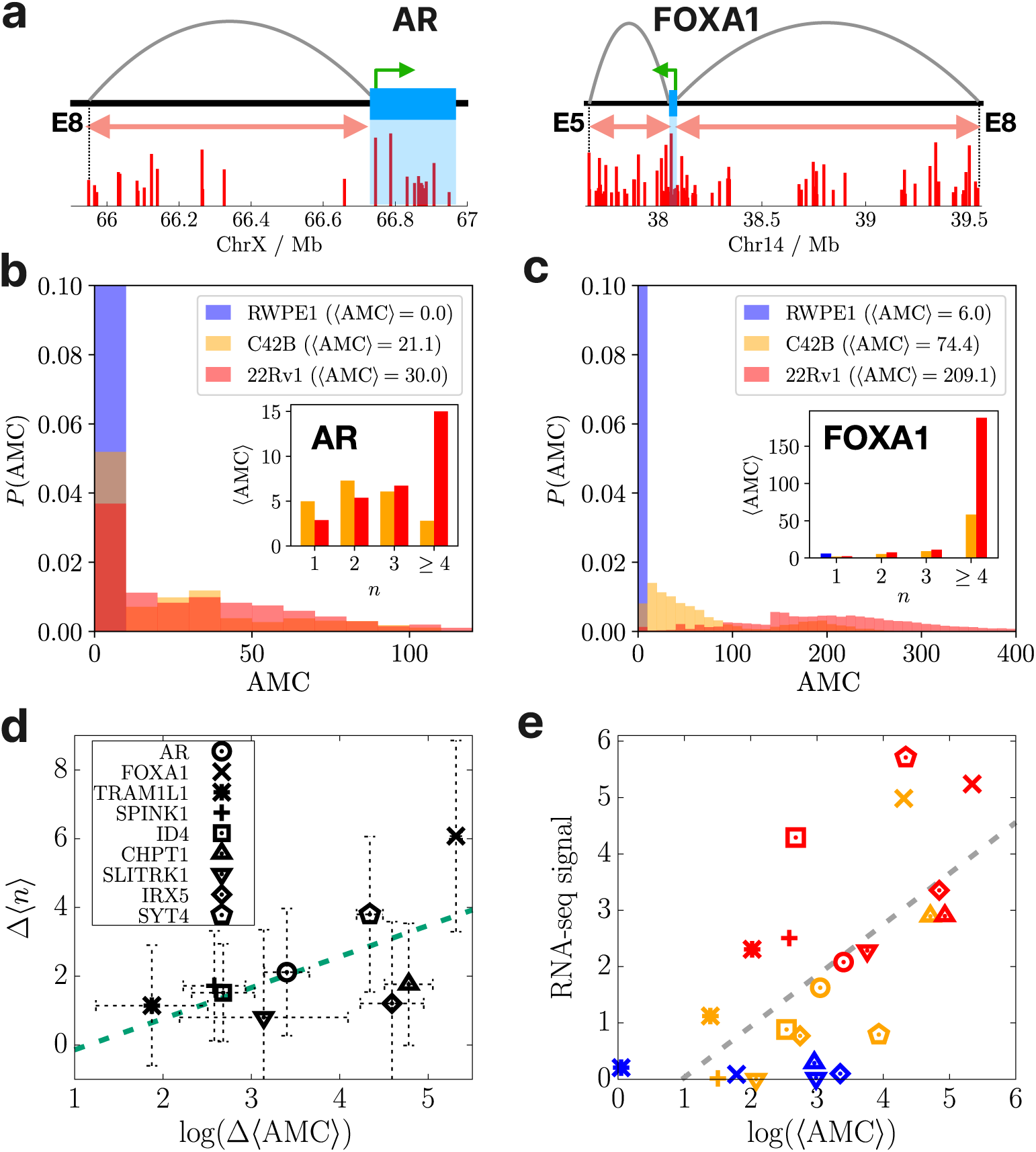
Activity-by-Multiway-Contact (AMC) model. (a) Schematic illustration for the range of enhancer candidates included in the AMC calculation. On the top panels, the blue rectangles are the gene bodies of *AR* (left) and *FOXA1* (right) in ChrX and Chr14, respectively. The bottom panel shows the ChIP-seq signals of H3K27ac for the gene regions in the 22Rv1 cell line. The green arrows indicate the direction of gene transcripts from the promoters. The gray arches are the most distant E–P interactions reported previously 4. The active H3K27ac peaks in the intervals covered by the red arrows, excluding the regions corresponding to the gene bodies (under blue shadows), are considered in the AMC calculations. (b) Probability distributions of the AMC score for the *AR* gene in different cell types. The inset shows the contribution of multiway contacts involving *n* enhancers to the mean AMC score for each cell line. (c) Same as panel b, except for *FOXA1* gene. (d) Increase in the average multiway contacts versus the logarithm of the difference between the mean AMC scores of 22Rv1 and RWPE1 for various genes. The green dashed line is the best linear fit (Pearson correlation coefficient (PCC) *r* = 0.63). (e) RNA-seq signals versus the logarithm of the mean AMC scores for the genes shown in panel d. The blue, orange, and red colors correspond to RWPE1, C42B, and 22Rv1. The gray dashed line is the best linear fit (PCC is *r* = 0.65).

To assess the broader applicability of the AMC model, we analyzed a comprehensive set of genes that show significant overexpression in 22Rv1 cell line. From the 22,236 genes with RNA-seq data [4], we identified 162 genes whose mean expression level in the 22Rv1 cancer cell line is at least 10-fold higher than that in the normal RWPE1 cell line. This stringent selection criterion ensures that we focus on genes with substantial cancer-associated over-expression. To ensure representative genomic coverage, we first selected one overexpressed gene from each chromosome, yielding a total of 22 genes (one per chromosome). Among these, 11 genes (*ADRA2A*, *CHPT1*, *SLITRK1*, *SYT4*, *KLK4*, *TRAM1L1*, *SPINK1*, *ID4*, *CLU*, *AR*, and *FOXA1*) have enhancer annotations available in the dataset of Rhie *et al.* [4], and thus their AMC scores were computed.

The results of the AMC scores for *AR* and *FOXA1* are shown in Figs. 5b and c. The distributions and mean values show that a large AMC score is obtained for the cancer cell lines, C42B and 22Rv1, in which the gene expression levels for *AR* and *FOXA1* are significantly enhanced relative to those in the normal cell line.

The AMC score also makes prediction on the contribution of multiway E–P contacts to the overall gene expression, which is not feasible based solely on pairwise contact probability. This is because the AMC score is computed for individual structures, naturally incorporating multiway contact information. Importantly, the AMC model allows us to separate contributions from 2-way, 3-way, and higher-order multiway contacts between promoters and enhancers. For the *AR* locus, the inset in Fig. 5b demonstrates that contacts involving more than 4 enhancers contribute most significantly to the AMC score in the 22Rv1 cancer cell line. This finding underscores the importance of multiple enhancers simultaneously interacting with promoters for *AR* gene in 22Rv1 cells, suggesting that the formation of complex regulatory hubs involving multiple enhancers is a key mechanism driving enhanced gene expression in prostate cancer. Similar observation is made for *FOXA1* as well (Fig. 5c). For the other nine genes selected, we observed similar patterns to *AR* and *FOXA1* (SI Figure 16). The increase in the average multiway contacts, Δ⟨*n*⟩ = ⟨*n*⟩_22Rv1_ − ⟨*n*⟩_RWPE1_, is positively correlated with the logarithm of Δ⟨AMC⟩ = ⟨AMC⟩_22Rv1_ − ⟨AMC⟩_RWPE1_ (Fig. 5(d)). Similarly, there is a positive correlation between the RNA-seq signals and the mean AMC scores for the genes in different cell types (Fig. 5e).

To account for intrinsic gene-specific effects that could obscure the correlation between AMC scores and gene expression, we computed RNA-seq – 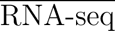 and ⟨AMC⟩ − 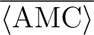 for each gene where the overline represents the average over the three cell lines for each gene. This analysis effectively removes gene-specific baseline effects, allowing us to focus on cell-type-specific variations. We analyzed 54 genes with enhancer information from the full 162 gene pool. The correlation coefficient *ρ* between RNA-seq 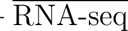 − and − ⟨AMC⟩ 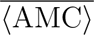 is 0.45, which is statistically significant (*p* = 8.2 × 10^−9^), demonstrating that AMC scores capture meaningful cell-type-specific regulatory changes beyond intrinsic gene expression differences (see SI Fig. 18).

### AMC model breast cancer progression

A recent study [33] reported high-resolution Micro-C maps for non-malignant MCF10A, pre-malignant MCF10AT1, and metastatic MCF10CA1a cells, together with changes in the gene expression. It was shown [33] that cancer progression was accompanied by changes in higher-order genome organization and by alterations in E–P contacts and enhancer activity at some progression-associated genes. Using Micro-C data, we applied the AMC model to assess if the connection between multiway E–P contacts and transcriptional changes holds in breast cancer progression. We focused on five genes which were particularly prominently changed in the MCF10 progression model, namely *SPRY1*, *SCNN1G*, *SCNN1B*, *COL12A1*, and *WNT5A*, which are highlighted in [33] (see SI for details).

*SPRY1* serves as a representative example (Fig. 6). Active enhancers were identified based on H3K27ac ChIP-seq (see SI for details) and 3D structures derived from 10kb-resolution Micro-C data using the HIPPS model (Fig. 6a,e). We quantified the enhancer promoter contacts as well as the AMC score. We find that the number of multiway E–P contacts in the *SPRY1* locus increases from the non-cancerous MCF10A state to the pre-malignant MCF10AT1 state and to the metastatic MCF10CA1a state (Fig. 6b). For the metastatic MCF10CA1a cell, the probability of *n* enhancers in contact with promoter *P* (*n*) is maximum at *n* = 3, in contrast to the MCF10AT1 and MCF10A cell lines. During the progression, the increases in RNA-seq expression is associated with increases of the mean AMC score (Fig. 6c). Thus, for this locus, stronger multiway E–P coupling is associated with increased transcription along the cancer progression.

**FIG. 6:**
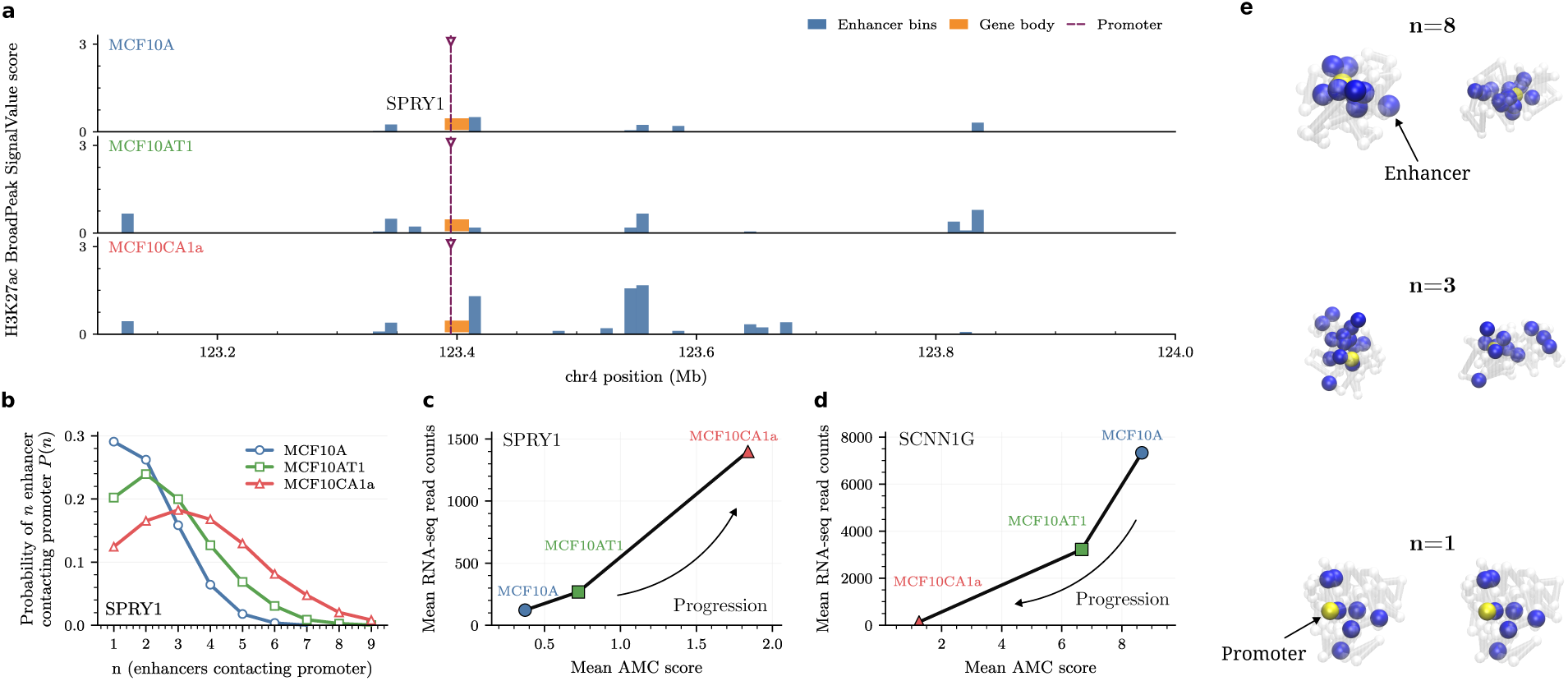
Multiway enhancer-promoter contact and expression analysis for *SPRY1* and *SCNN1G* across breast cancer progression. **(a)** Linear genomic track across the *SPRY1* locus showing the H3K27ac broadPeak score in MCF10A (non-cancerous), MCF10AT1 (pre-malignant), and MCF10CA1a (metastatic). Putative enhancer bins with the gene body and promoter position. The bin size is 10kb. **(b)** Distribution of enhancer-promoter (E–P) contacts, showing the probability of*n* number of enhancers in contact with promoter for each cell line. **(c)** Mean RNA-seq read counts versus mean AMC score for *SPRY1* in MCF10A, MCF10AT1, and MCF10CA1a. Increased mean AMC score are accompanied by increased *SPRY1* expression in the three cell lines. **(d)** Same as **(c)** for gene *SCNN1G*. Decrease of RNA-seq counts is accompanied by decreased mean AMC score. **(e)** Examples of chromatin conformation for *SPRY1* gene region for different number of enhancer in contact with promoter *n*. The enhancers and promoter are shown in blue and yellow, respectively. The other loci are in semitransparent gray. The analysis shown in this figure is detailed in SI.

For *SCNN1G* (Fig. 6d and SI Fig. 20), the RNA-seq level decreases during the progression and is accompanied by a progressive decrease in the mean AMC score (Fig. 6d). The results for the remaining genes are shown in SI Figs. 19–23. *SCNN1B* and *COL12A1* follow qualitatively the same trend, namely, changes in RNA-seq are positively correlated with the changes in multiway E–P contacts and the AMC score. *WNT5A* is an exception. For *WNT5A*, the RNA-seq level increases strongly from MCF10A to MCF10AT1 and then de-creases to a low level in MCF10CA1a, whereas both the mean number of E–P contacts and the mean AMC score decrease along the progression. These examples show that changes in the multiway E–P contacts, predicted by the HIPPS model, when combined with biochemical activity explains the locus-dependent activation (*SPRY1*, *COL12A1*) and repression (*SCNN1G*, *SCNN1B*) during breast cancer progression with exception of the *WNT5A*.

## III. DISCUSSION

We developed a computational framework that combines the multiway interactions (calculated using experimental contact maps) between enhancers and promoters and ChIP-seq data to explain the origin of the increased expression of key genes in prostate cancer cells relative to normal epithelium. Remarkably, the only input in our computations is the readily available two-dimensional Hi-C data for normal and cancer cell lines. The three-dimensional coordinates of chromosomes, calculated using the Hi-C data, permitted us to calculate a variety of structural properties, that proved to be the starting point in understanding the variations in gene expression across the cell lines. Let us first summarize the major findings of our investigation. (1) The enhancer–promoter distance distributions in the *AR* and *FOXA1* genes are broad. In the *AR* gene, the mean E–P distances decrease in the cancer cells, with respect to the normal epithelium. In contrast, they are roughly constant or in-crease modestly in *FOXA1* (Fig. 2). It follows that models requiring E–P contacts cannot explain the increase in gene expression as the normal cells become cancerous. (2) The number and importance of multiway contacts increase in cancer cells for the *AR* gene (Fig. 3). In contrast, the probability of multiway contact formation is nearly the same in the 22Rv1 cancer cell as it is in the normal RWPE1 cell (Fig. 4). These results imply that neither multiway contacts nor E–P distances can account for enhanced gene expression of *AR* and *FOXA1* genes. (3) The AMC model that combines multiway contacts and ChIP-seq signals explains the similar extent of gene expression in *AR* locus and *FOXA1* (Figs. 5 **b** and **c**). We also show that the predictions of the model correlate with the RNA-seq data for several other genes (Figs. 5 **d** and **e**). Because the outcome of the AMC model is the distribution of the AMC scores (Figs. 5 **b** and **c**), single cell experiments [41] are required to validate the predictions.

### On spatial proximity between enhancers and promoters

Although appealing, the model requiring the E–P distances should be close enough for direct physical contact between enhancers and promoters for gene activation, is not always supported by experiments [18, 42]. Active enhancers and their target promoters often maintain surprisingly large distances even during transcriptional activation [43, 44] typically a few hundred nanometers, which are too large for physically establishing contact. For instance, in *Drosophila* embryos, the average E–P distance during active transcription was ⟨*R*⟩ ∼ 280 nm with a standard deviation *σ* ∼ 120 nm (⟨*R*⟩*/σ*=2.3) [42]. Intriguingly, some studies even observed an increase in E–P distance upon transcriptional activation, as is the case with the *SHH* promoter and the associated enhancer [45]. These findings suggest that direct E–P contact is not necessary for gene activation. The cited experiments have investigated gene activation by a single enhancer whereas our study shows that the same conclusion holds when multiple enhancers are involved. In particular, ⟨*R*⟩*/σ* ≈ is in the range of 2.3 ∼ 2.5 (Figs. 2b,d). Finally, the difficulty of directly correlating contact frequency (inferred from chromatin conformation capture experiments) and spatial E–P proximity using imaging experiments [46–50], due to chromatin conformational heterogeneity, makes conclusions based on proximity measurements alone tenuous.

A model that explains the activity of enhancers without the need for close physical proximity is based on the formation of phase-separated condensates [51]. In this model, enhancers act as nucleation sites for membraneless protein condensates that arise through weak, multivalent interactions via intrinsically disordered regions, often found in transcription factors. Because these condensates can span several hundred nanometers, stable, direct contacts between an enhancer and its target gene is not strictly required and enhancer proximity to a target promoter is not essential for modulating transcriptional bursting. Interestingly in the context of our observations on multi-enhancer effects, the combined action of multiple enhancers is thought to promote the formation of transcriptional condensates [51].

### Importance of multiway contacts

The changes in E–P distances and multiway contacts provide a coherent picture of chromatin reorganization during prostate and breast cancer progression. In the *AR* locus, enhancers and promoters move closer in 3D space, especially in the 22Rv1 cell line, consistent with the observed upregulation of *AR*. Multi-way E–P contacts also increase in cancer cells, indicating the formation of regulatory hubs that bring multiple enhancers into proximity with the promoter. Enhancers E4–E7 play particularly prominent roles in these higher-order interactions. Importantly, the ensemble-averaged number of enhancers in contact with the promoter also correlates with the increase in gene expression, found in breast cancer progression (Fig. 6). The E–P contact structure in MCF10A (non-cancerous), MCF10AT1 (pre-malignant), and MCF10CA1a (metastatic) cell lines is consistent with the changes in gene expression along the progression.

In contrast, the *FOXA1* locus exhibits minimal changes in the pairwise E–P distances despite a comparable enhanced overexpression as in the *AR* locus. Just like the *AR* locus, *FOXA1* also shows strong multiway contact patterns, suggesting that simultaneous cooperative interactions between several enhancers drive transcription. However, in *FOXA1* the multiway contacts in the normal and cancer cells are equally important whereas in *AR* locus triple or higher order contacts take on greater importance as the cells become cancerous. From these findings we surmise that changes in chromatin structures alone, no matter how detailed, are inadequate to link to gene expression changes.

The AMC analysis further integrates epigenetic activity and multiway contacts, revealing that both *AR* and *FOXA1* have higher AMC scores in cancer cells, correlating with increased expression (Fig. 5). Across several genes, the AMC score positively correlates with both multiway contact frequency and RNA expression, highlighting that enhancer activity combined with three-dimensional chromatin structures could explain transcriptional output better than distance alone. Together, these findings suggest that gene activation in prostate cancer arises from the combined effects of compartment switching, increased E–P proximity, and formation of multi-enhancer regulatory hubs [25].

### Promoter-Enhancer Hubs

Several recent experiments, using optical imaging [3, 25, 52–55] and single-cell Micro-C [56], suggest that multiple enhancers and the associated promoter spatially colocalize (not necessarily in contact) producing promoter-enhancer hubs (PEHs). However, the probability of forming such hub-like structures is low in individual cells (Fig. 3) because it requires more than two enhancers be simultaneously within physically reasonable distance to the promoter even if they are separated by a large genomic distance. In the PEH model, enhancers, promoters, and possibly other regulatory elements are localized in a small region in the nucleus. As a result, the concentration of the relevant species required for transcription are confined to a small volume, which would facilitate gene regulation. The model does not require highly specific E–P interactions. We ought to emphasize that the emergence of PEHs is a natural consequence of the massively heterogeneous organization of chromatin [47, 48, 57]. By concentrating the various components in small volume, the rate of transcription is likely to increase substantially. Moreover, the PEH model qualitatively explains how multiple enhancers can simultaneously regulate genes by bringing them into the small hub [51, 58–62]. Our calculations show that three enhancers make contacts with the promoter simultaneously (probability on the order of (10 - 15)%). Although this may be an overestimate considering that the two enhancer alleles in an individual nucleus may differ in their activity status, which is not captured in our analysis. Hubs consisting of more than four enhancers are possible (see Fig. 4 in [56]), but their probability decreases (Figs. 3 and 4) as the number of multiway interactions increases.

Our finding that three and higher order E–P interactions are prevalent for over-expressed genes in prostate and breast cancer cells suggests that E–P hubs could be common in other cancers as well, as pointed out recently. Indeed, recent studies [25, 53] showed that E–P hubs involving three way contacts between *SOX9* and *MYC* alleles and promoters form hubs during transcription, albeit with low probability. Our calculations show that the probability of forming three enhancers in contact with the promoter in both prostate and breast cancer cell is small (Fig. 3 and Fig. 6), in accord with experiments.

### Limitations

Our study has several limitations. First, the analysis of prostate cancer is based on 40 kb resolution Hi-C data, which could obscure important structural features. Each locus on this scale may include multiple enhancers, limiting our ability to resolve fine-grained interactions. Further studies using higher-resolution data such as Micro-C [33, 63] or RCMC [64] could reveal more precise structural details and test the robustness of the AMC model. In this regard, the use of single cell Micro-C to shed light on the importance of multi-way E–P contacts in cancer cells would be most welcome. Second, our definition of enhancer activity relies solely on H3K27ac as a proxy, which does not fully capture the complexity of enhancer states. Incorporating additional epigenetic marks such as H3K4me1, H3K36me3, CTCF binding, chromatin accessibility (ATAC-seq) as well as potential cooperativity between the activities will allow a more comprehensive understanding of E–P regulation and the interplay between the three-dimensional genome architecture and epigenetic landscapes.

## IV. METHODS

### Generation of 3D structures from Hi-C and Micro-C data using HIPPS

To generate the ensemble of chromosome structures from the experimentally determined population averaged Hi-C map, we used the HIPPS method [28], which is conceptually analogous to generative modeling in machine learning [65]. The objective is to derive the joint distribution function of the variables, ***x****_i_*’s, specifying the three-dimensional coordinates of the chromatin loci. The input needed is the pairwise contact probability/frequency matrix that is obtained from bulk Hi-C/Micro-C experiments.

There are two main steps in the implementation of HIPPS.

1. We first convert the Hi-C contact frequency matrix to the average distance matrix using the power-law relation between the contact probability, ⟨*P_ij_*⟩, and the mean spatial distance ⟨*r_ij_*⟩. For standard polymer models, such as Rouse [66] or the Generalized Rouse Model (GRMC) [48, 67], the power-law scaling can be analytically derived. For chromosomes, a similar empirical relation holds [27, 48, 68, 69],

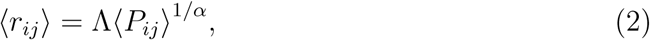

although the exponent *α* differs from standard polymer models. Experiments [68, 70] and simulations [26] have suggested that *α* ≈ 4. We have previously used HIPPS with *α* ≈ 4 to accurately reproduce the structures of chromosomes in the interphase [28] and the changes that occur during cell cycles [71]. The Λ value in Eq. 2 was set to the threshold contact distance, *r_c_* = 1.5*σ*. For a coarse-grained locus, representing 40 kb of chromatin, *σ* = 160 nm is the effective diameter.

2. To generate a 3D ensemble of chromatin conformations, we computed the joint probability distribution of locus coordinates *x_i_* subject to the constraints calculated from Hi-C contact map,

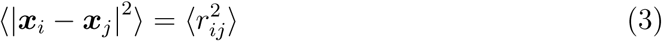

for all loci pairs (*i, j*). The maximum entropy principle yields the joint distribution,

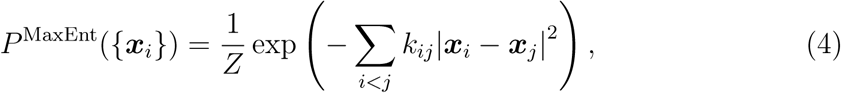

where *Z* is the normalization factor and *k_ij_*’s are the Lagrange multipliers that enforce the constraint in Eq.3. The values of *k_ij_* were determined using an iterative scaling algorithm [28, 72]. One of the advantages of Eq. 4 is that it can be sampled efficiently because it is a multivariate normal distribution.

By using the two steps described above, an ensemble of 3D chromosome structures that are consistent with the Hi-C map is readily generated. Given that typical topologically associating domains (TADs) range from 0.5 to 3 Mb in size [22, 73], we reconstructed chromatin structures using 10 Mb segments with 1 Mb overlaps, enabling us to capture higher-order organization across multiple TADs. Since our analysis focuses on enhancer–promoter (E–P) interactions in normal and cancer cells, the 10 Mb genomic window provides sufficient structural detail.

## Data availability

Hi-C datasets at a 40 kb resolution from Rhie et al. [4] (accession number GSE118629) were used for the normal prostate epithelial cell line RWPE1 and the prostate cancer cell lines C42B and 22Rv1. For the prostate cancer cell line MDAPCa2b, we employed the Hi-C data at the same resolution from Rebeca et al. [5] (accession number GSE172099). Each Hi-C contact map was independently normalized on a per-chromosome basis prior to analysis with HIPPS. The ChIP-seq datasets targeting H3K27ac for 22Rv1 (ENCSR391NPE), C42B (ENCSR279KIX), and RWPE1 (ENCSR203KEU) were retrieved from the ENCODE database. The corresponding narrowPeak files were processed and normalized to counts per million (CPM) to ensure compatibility of signal intensity across the cell lines. The Micro-C datasets used for MCF10A, MCF10AT1 and MCF10CA1a are available in the GEO repository with accession GSE254045. H3K27ac data for MCF10A, MCF10AT1 and MCF10CA1a are obtained from the GEO accession GSE229295. RNA-seq for MCF10A, MCF10AT1 and MCF10CA1a are obtained from the GEO accession GSE320216.

## Code availability

The code for implementing the latest version of HIPPS model used in this work with detailed user instructions is available at the GitHub repository: https://github.com/anyuzx/HIPPS-DIMES.

## Acknowledgments

This research was supported by the National Science Foundation (PHY 2310639) and the Collie-Welch Chair through the Welch Foundation (F-0019). S.S. acknowledges that this work was supported by Sungshin Women’s University Research Grant of H20250122. This research was supported by the Intramural Research Program of the National Institutes of Health (NIH), National Cancer Institute NCI, Center for Cancer Research through grant 1-ZIA-BC010309 to T.M. The contributions of the NIH author(s) were made as part of their official duties as NIH federal employees, are in compliance with agency policy requirements, and are considered Works of the United States Government. However, the findings and conclusions presented in this paper are those of the author(s) and do not necessarily reflect the views of the NIH or the U.S. Department of Health and Human Services.

