## Supplementary Information for "Connecting multiway enhancer-promoter interactions to changes in gene expression in cancer"

**Chromatin A/B compartments from Hi-C contact maps**

We identify the A (active euchromatin) and B (inactive heterochromatin) compartments using the principal component analysis (PCA). Starting from the raw contact matrix $C_{ij}$, we constructed an observed/expected (O/E) normalized matrix $C^*_{ij}$ by dividing each contact frequency by the expected contact frequency at the same genomic separation $|i-j|$. The expected frequency is computed as the average contact count over all pairs of loci separated by $|i-j|$ along the chromosome. Specifically, we define

$$C^*_{ij} = \frac{C_{ij}}{\sum_{k=1}^{N}\sum_{l\geq k+1}^{N} C_{kl}\delta_{|k-l|,|i-j|}/(N-(|i-j|))}. \quad (1)$$

We converted $C^*_{ij}$ into a Pearson correlation matrix $\rho_{ij}$ by calculating the correlation between row vectors $C^*_i$ and $C^*_j$,

$$\rho_{ij} = \frac{\mathrm{cov(C^*_i C^*_j)}}{\sqrt{(\mathrm{cov(C^*_i C^*_i)cov(C^*_j C^*_j)})}} \quad (2)$$

where $\mathrm{cov}(C^*_i, C^*_j)$ denotes the covariance between the normalized vectors $C^*_i$ and $C^*_j$.

Subsequently, we performed principal component analysis (PCA) on the Pearson correlation matrix $\rho_{ij}$. The first principal component (PC1), given by the leading eigenvector, provides a scalar score for each locus. Loci with opposite PC1 signs are assigned to different compartments. To map the PC1 sign to the A (active) or B (inactive) compartment, we compared PC1 with epigenomic tracks from reference human genomes, using active histone marks (e.g., H3K27ac) as a guide [1, 2]. The sign enriched for active marks correspond to the A compartment, and the one with the opposite sign is identified as the B compartment [3, 4]. The changes in the compartment structure based on the PCA analysis across the cell lines for chromosome X shows (Fig. SI-1) an increase in the A compartments as the normal epithelium (RWPE1) becomes cancerous. The changes in the the compartment structure provides a qualitative explanation of the enhanced gene expression in the cancer cells.

* These authors contributed equally.
†

**Clustering**

We determined the three-dimensional ensemble of conformations using the HIPPS method [5]. To discern the range of conformations sampled by the chromosomes in the different cell lines, we grouped them into clusters based on a similarity measure. Conformations within a cluster are expected to be similar. Similarity of conformations is assessed using the distance metric, $D_{mn}$,

$$D_{mn} = \sqrt{\frac{1}{N^2} \sum_{i,j} \left( r_{ij}^{(m)} - r_{ij}^{(n)} \right)^2}. \tag{3}$$

In the above equation, $r_{ij}^{(m)}$ and $r_{ij}^{(n)}$ are the Euclidean distances between the $i^{th}$ and $j^{th}$ loci in conformation $m$ and $n$, respectively. We computed the distance metric (Eq. 3) after structure alignment to remove translational and rotational degrees of freedom. First, each conformation was translated so that its center of mass coincided with that of a reference conformation. We then rotated each conformation to minimize the root-mean-square deviation (RMSD) from the reference. This procedure ensures that the distance metric reflects intrinsic structural differences rather than artifacts arising from overall translation or rotation.

We analyzed a dataset of 10,000 conformations generated with the HIPPS method [5], which is used to construct a 10,000 $\times$ 10,000 distance matrix. Each entry, $D_{mn}$, quantifies the dissimilarity between conformations $m$ and $n$ as defined in Eq. 3. We then applied agglomerative hierarchical clustering to group conformations into structurally similar clusters based on these pairwise distances. To select the number of clusters, we evaluated the silhouette score [6, 7], where larger values indicate better within-cluster cohesion and stronger separation between clusters (Fig. SI-2a).

To visualize the clustering results, we used t-distributed stochastic neighbor embedding (t-SNE) [8] to embed the conformations into two dimensions based on the distance matrix. This low-dimensional representation, which provides an intuitive view of how conformations and clusters are arranged relative to one another (Fig. SI-2b), shows that the chromatic structures partition into two disjoint clusters. There is little mixing between the 3D structures belonging to the clusters.

### Locus size and contact threshold

The resolution of the Hi-C maps the prostate cancer is 40 kb [9]. To determine the size of each locus, we assume that the radius of gyration of the 40 kb chromatin polymer, $R_g(40\,\mathrm{kb})$, scales as $N^{1/3}$, corresponding to a fractal globule. Using the estimate [10] that 1.2kb locus size is $\approx 50\,\mathrm{nm}$, the size of a locus, $\sigma$, at 40kb is,

$$\sigma(40\,\mathrm{kb}) = \left(\frac{40\,\mathrm{kb}}{1.2\,\mathrm{kb}}\right)^{0.33} 50\,\mathrm{nm} \approx 160\,\mathrm{nm}. \tag{4}$$

A contact, $C_{ij}$, in a given conformation is defined as,

$$C_{ij} = \begin{cases} 1, & \text{if } r_{ij} \leq r_c \\ 0, & \text{otherwise.} \end{cases} \tag{5}$$

The mean value of the number of contacts (contact probability) is obtained by averaging over all the conformations and is denoted as $P_{ij} = \langle C_{ij} \rangle$. To determine the optimal value of $r_c$, we calculated the contact map (see Eq. (3) in the main text) by varying $r_c$ (Fig. SI-3). In the range $\sigma \leq r_c \leq 1.5\sigma$, the agreement between experimental and the calculated contact maps are in excellent agreement (Fig. SI-3B). This also reflected in the Pearson correlation coefficient (Fig. SI-3C) which exceeds 0.95. Accordingly, we set $r_c = 1.5\sigma = 240\,nm$, which is less than the typical mean distances between an enhancer and the promoters, AR and FOXA1 (see Fig. (2) in the main text).

### Jensen-Shannon Divergence (JSD)

The calculated distribution functions (see below) are compared with each other to assess the extent to which they overlap. The symmetric Jensen–Shannon divergence (JSD), derived from the Kullback–Leibler (KL) divergence, provides a measure of the similarity between two probability distributions. The value of JSD is zero when the two distributions are identical. We define JSD between two distributions $p_1$ and $p_2$ using,

$$\mathrm{JSD}(p_1||p_2) = \frac{1}{2}\big(\mathrm{KL}(p_1||M) + \mathrm{KL}(p_2||M)\big), \tag{6}$$

where $M = (p_1 + p_2)/2$ is the mean distribution. The Kullback–Leibler divergence is,

$$\mathrm{KL}(p_1||M) = \sum_i p_1(i) \log \frac{p_1(i)}{M(i)}, \tag{7}$$

where the summation is over all the possible values of $i$. In the above equation, $p_{1/2}(i)$ and $M(i)$ represent the probabilities of $i$ under the distributions $p_{1/2}$ and $M$, respectively.

### Structural features

**Sizes and shapes:** Using the ensemble of 3D conformations, we calculated the distribution of the radius of gyration, $R_g$, and the shape parameters, $\kappa^2$ and $S$. The latter two measures are useful descriptors in characterizing polymer shapes [11–13]. All three quantities are calculated using the inertia tensor, $T_{\alpha\beta}$,

$$T_{\alpha\beta} = \frac{1}{2N^2} \sum_{i,j=1}^{N} (r_{i\alpha} - r_{j\alpha})(r_{i\beta} - r_{j\beta}), \tag{8}$$

where $N$ is the number of loci, $r_{i\alpha}$ is the $\alpha^{th}$ component of the position of the locus $i$ and $\alpha\beta = x, y, z$ [13, 14]. The radius of gyration is given by,

$$R_g^2 = \lambda_1 + \lambda_2 + \lambda_3 \tag{9}$$

where $\lambda_1$, $\lambda_2$ and $\lambda_2$ are the eigenvalues of $T_{\alpha\beta}$. The shape anisotropy ($\kappa^2$), with bounds $\kappa^2$ is $0 \le \kappa^2 \le 1$, is defined as,

$$\kappa^2 = \frac{3}{2} \frac{\lambda_1^2 + \lambda_2^2 + \lambda_3^2}{(\lambda_1 + \lambda_2 + \lambda_3)^2} - \frac{1}{2} \tag{10}$$

For a sphere $\kappa^2$ is zero and for a rod it is unity. The overall shape $S$ is,

$$S = \prod_{i=1}^{3} \frac{\lambda_i - \overline{\lambda}}{\overline{\lambda}}, \tag{11}$$

$\overline{\lambda} = \frac{\lambda_1+\lambda_2+\lambda_3}{3}$. For a sphere the value of $S = 0$. A positive (negative) value of $S$ implies that the conformation is prolate (oblate) ellipsoid.

The distributions of $R_g$ in all the cells lines are roughly similar as are their average values

((Fig. SI-4a). The anisotropy ($\kappa^2$) and shape $S$ are quantitative descriptors of the extent to which chromatin conformations deviate from a sphere. In the RWPE1, the average $\kappa^2$ for cluster-1 is 0.17, with a standard deviation of 0.10, indicating that the structures are symmetric and roughly sphere-like (for a perfect sphere $\kappa^2 = 0$) (Fig. SI-4b). In contrast, the distributions of $\kappa^2$ in Cluster-2 conformations are broad in all the cell lines with the mean values $\kappa^2 \approx 0.26$, suggesting increased anisotropy. Interestingly, the average values of $\kappa^2$ in the two clusters do not change significantly across the cell lines. Similarly, the distributions of the shape parameter, $S$, in cluster-1 are similar across all the cell lines (Fig. SI-4c). However, there is a greater spread in the average values of $S$ (the mean varies from 0.18 to 0.34). The average values of $S$ in all the cell lines are positive, which implies that the chromosome conformations resemble prolate ellipsoids. The degree of prolateness is greater in cluster-2 structures (Fig. SI-4).

**Angle between the promoter and enhancers:** We also calculated the angles between the enhancers using the promoter as an anchor point change in the different cell lines. Fig. SI-5(a) presents a schematic representation of the angle $\theta_{E4,E6}$, defined by the vectors connecting P-E4 and P-E6. The angles vary modestly in cancer cell lines compared to the normal cell line. For instance, the $\langle\theta_{E6,E4}\rangle$ (in degrees) is 41 with the standard deviation of 29. In the cancer cell lines MDAPCa2b, C42B, and 22Rv1, the angle changes to 39, 43 and 45, respectively. Similarly, $\langle\theta_{E6,E5}\rangle$ also shows only slight increase in the angle for the cancer cell lines compared to the normal cell line (see Fig. SI-5(b)). To illustrate the relation between the angle ($\theta_{E6,E5}$) and $R_{P,E6}$, we computed the contour plot between these quantities (Fig. SI-5(c)). In contrast to the present findings, a recent imaging study [15] found that the angle made by labeled enhancers of the *PGR* gene, the progesterone receptor gene in breast decreases by about thirty degrees upon estradiol (a hormone that regulates transcription in mammalian cells) signaling even though the genomic location of the enhancers are comparable to those in the AR locus. It is likely that the orientation of enhancers with respect to the promoter would depend on the cell lines and the chromatin.

**Conformational heterogeneity in the AR locus is cell type specific**

Because a given contact exists only in a small fraction of population of cells, the chromosome conformations vary from cell to cell [16, 17], a feature that is masked in ensemble averages. Consider the single-cell distance map using 3D coordinates for each conformation in the two clusters for the region containing the AR gene in the four cell lines. In RWPE1, the loci in cluster-1 are in proximity (SI Fig. SI-8a), including those that are far from the diagonal, implying that the structures are compact. In contrast, the smaller single-cell distances in the structures in cluster-2 are predominantly between adjacent loci. The distances between the loci that are further from the diagonal are greater. The single-cell images, displayed below the DMs in SI Fig. SI-8a, show that the structures from both the clusters are substantially more heterogeneous than could be anticipated from averages. Similar trends are found in the three cancer cell lines. The structures associated with the conformations in cluster-1 are more compact than the ones in cluster-2. There are substantial cell-to-cell variations in the conformations in both the clusters, which reinforces the general notion that the chromatin conformations are massively heterogeneous [16].

To quantitatively assess the differences between the conformations in the different cell lines, we calculated the distribution of the radius of gyration ($R_g$) for each cluster separately. From the results in Fig. SI-8b a few conclusions can be drawn. (1) The mean radii of gyration ($\langle R_g \rangle$s) the in the two clusters are roughly independent of the cell lines. The $\langle R_g \rangle$ values of the in cluster-1 (cluster-2) are $\approx (0.29 \pm .03)\mu m$ ($\approx (0.39 \pm .03)\mu m$). (2) The $R_g$ distributions are broader in cluster-2 than in cluster-1 (Fig. SI-8b). Importantly, the $R_g$ distributions of conformations belonging to cluster-1 (blue) do not depend on the cell line. The Jensen-Shannon divergence (JSD) values between the three cell lines, with respect to the RWPE1, are roughly similar (JSD values are utmost 0.10) for cluster-1 conformations. There is a greater spread in the JSD values between the RWPE1 and the three cancer lines for $R_g$ distributions (orange) from cluster-2 conformations. (3) Despite the similarities, there is a notable difference. The size of cluster-2 in the cancer cell lines is larger compared to the normal epithelium. Clearly, there is a shift in the redistribution of structures between the clusters, which is a reflection of the underlying changes in the chromatin organization that occurs in the transition from normal to cancer cells. The observed changes are a reflection of the alteration of in the epigenetic states. For analyses of shape parameters ($\kappa^2$ and $S$) for the chromatin conformations see Fig. SI-4.

**Enhancer-promoter distance distribution functions**

In the main text the distribution of distances between two enhancers and promoter for the AR and FOXA1 are presented (Fig. 2 in the main text). For completeness the distributions of E-P distances for all the eight promoters are for both these genes are shown in Figs. SI-6 and SI-7. We examined additional regions on different chromosomes that house the genes FOXA1, RBM11, and SHH to assess if mean distances are indicators altered altered gene expression ib cancer cells.

**FOXA1** In prostate cancer cells, FOXA1 plays an important role in gene regulation. Many overexpressed genes in these cells have the FOXA1 motif in their promoter regions, establishing its involvement in activating the gene. FOXA1 itself is significantly overexpressed in prostate cancer cell lines, showing a a substantial fold increase in 22Rv1 and 56 in C42B, compared to normal cell lines [9].

We determined the 3D structures of the region containing the FOXA1 locus by generating 10,000 conformations using HIPPS for the segment of chromosome 14 spanning from 36Mb to 44Mb. Fig SI-7(A) compares the contact probability map from the HIPPS generated structure with the experimental Hi-C contacts. A zoomed-in view focusing on the FOXA1 region is presented in Fig SI-7(B). The lower panel illustrates the location of the FOXA1 promoter and the associated enhancers. FOXA1 is regulated by eight enhancers located that are located both upstream and downstream of the promoter unlike the AR locus in which all the enhancers are upstream of the gene. In Fig SI-7(C), we present the differences in the distance maps between the normal RWPE1 and the cancer cell lines C42B and 22Rv1.

The calculated distance distribution of the enhancer and promoter ( Fig SI-9 and Fig SI-10) revealed a clear trend: for all the enhancers, with the exception of E1, the average spatial distance in the cancer cell line is consistently shorter compared to the normal cell line. This suggests that chromatin structure is likely to be more compact in cancerous cells, potentially influencing gene regulation mechanisms. For the enhancers that are genomically farther from the promoter, the disparity in their spatial distances between cancer and normal cell lines increases. In other words, the distal enhancers show the largest structural differences. Notably, enhancers E7 and E8 exhibit the most significant divergence in spatial proximity when comparing normal and cancerous cells, suggesting a stronger disruption in chromatin organization at these sites.

**RBM11:** We extended the analysis by focusing on an intriguing gene RBM11 gene located on chromosome 21. The promoter of RBM11 is positioned at approximately 15.5 Mb on chr21. Unlike AR locus and FOXA1, RBM11 is regulated by a "single" enhancer situated several TADs away on the same chromosome (Fig SI-11(A)). We first calculated the average distance between the promoter and all other loci of the given region (Fig SI-11(B)). The distance enhancer-promoter distribution (Fig SI-11(C)) shows that the peak position decreases in the MDAPCa2b cell line by about 70 nm relative to RWPE1 but increases by 140 nm and 90 nm in C42B and 22RV1, respectively. The widths of the distance distributions are roughly similar. There are substantial differences in the distributions between the four cell lines and those in the two control cases (Fig SI-11(C)). The distance changes are not always good indicators of enhancement in gene expression in cancer cells.

**SHH:** The enhancer-promoter distance for the SHH gene, in contrast to the AR gene, increases upon gene activation. Using 3D-FISH and 5-C techniques, it was demonstrated [18] that in neural progenitor cells (NPCs), where SHH is active, the enhancer-promoter distance is greater than in the repressed state observed in mouse embryonic stem cells (mESCs). We used the 5C contact data to generate 10,000 structures using HIPPS. We simulated a 1.4Mb region of chr5 spanning from 28.6MB to 30.0Mb, as presented in Fig. SI-12A. This region contains the promoter and the 4 enhancers (SBE6, SBE2/3, SBE4, and SBE5) associated with the SHH gene. The average distance between the SHH promoter and the enhancers computed from the distance distribution (see Fig. SI-12B and C) shows modest change in the mean distance between the enhancer and promoters.

### Activity-by-multiway-contact (AMC) model

The Activity-by-Multiway-Contact (AMC) model extends the Activity-by-Contact (ABC) framework [19] by defining a gene-level score that integrates enhancer biochemical activity with 3D chromatin structure at the level of individual conformations. The central idea is that the regulatory input to a promoter in a given conformation is the sum of the activities of all enhancers that are simultaneously in spatial contact with that promoter. Unlike the ABC score, which is defined per enhancer using ensemble-averaged contact probabilities from Hi-C, the AMC score is computed for each 3D conformation and thus naturally incorporates multiway (simultaneous) enhancer–promoter contacts.

Let $g$ denote a gene, $\mathrm{P}(g)$ the associated promoter, E an enhancer, and S the set of enhancer candidates within the analysis window for the gene. For a chromatin conformation $\xi = \{\boldsymbol{r}_1, \boldsymbol{r}_2, \ldots, \boldsymbol{r}_N\}$, the contact indicator between enhancer E and the promoter $\mathrm{P}(g)$ is

$$C_{\mathrm{E},\mathrm{P}(g)}(\xi) = \mathbf{1}\big(|\boldsymbol{r}_{\mathrm{E}} - \boldsymbol{r}_{\mathrm{P}(g)}| < r_c\big)\,, \tag{12}$$

where $\mathbf{1}(\cdot)$ equals unity if enhancer E is within the cutoff distance $r_c$ from the promoter $\mathrm{P}(g)$, and zero otherwise. The number of enhancers that are simultaneously in contact with the promoter $\mathrm{P}(g)$ in conformation $\xi$ is $n_{\mathrm{P}(g)}(\xi) = \sum_{\mathrm{E}\in\mathrm{S}} C_{\mathrm{E},\mathrm{P}(g)}(\xi)$. The AMC score for gene $g$ in conformation $\xi$ is

$$\mathrm{AMC}(g;\xi) = \sum_{\mathrm{E}\in\mathrm{S}} A_{\mathrm{E}}\, C_{\mathrm{E},\mathrm{P}(g)}(\xi), \tag{13}$$

where $A_{\mathrm{E}}$ is the biochemical activity of enhancer E, quantified here by H3K27ac ChIP-seq read counts (normalized to counts per million). If activities of all the enhancers are equal, $\mathrm{AMC}(g;\xi)$ reduces to $n_{\mathrm{P}(g)}(\xi)$.

The AMC score can be decomposed into contributions from conformations with exactly $m$ enhancers in contact with the promoter:

$$\mathrm{AMC}(g;\xi) = \sum_{m=1}^{|\mathrm{S}|} \delta_{n_{\mathrm{P}(g)}(\xi),m} \sum_{\mathrm{E}\in\mathrm{S}} A_{\mathrm{E}}\, C_{\mathrm{E},\mathrm{P}(g)}(\xi) = \mathrm{AMC}^{(1)}(g;\xi) + \mathrm{AMC}^{(2)}(g;\xi) + \cdots + \mathrm{AMC}^{(|\mathrm{S}|)}(g;\xi), \tag{14}$$

with $\mathrm{AMC}^{(m)}(g;\xi) = \delta_{n_{\mathrm{P}(g)}(\xi),m} \sum_{\mathrm{E}\in\mathrm{S}} A_{\mathrm{E}}\, C_{\mathrm{E},\mathrm{P}(g)}(\xi)$, where $\delta_{n,m}$ is the Kronecker delta. The term $\mathrm{AMC}^{(m)}$ is nonzero only when precisely $m$ enhancers are in contact with the promoter, and in that case it equals the sum of the activities of those $m$ enhancers. This decomposition quantifies the contribution of 1-way, 2-way, 3-way, and higher-order multiway contacts to the AMC score in each conformation.

Averaging Eq. 13 over the conformational ensemble yields $\langle \mathrm{AMC}(g) \rangle = \sum_{\mathrm{E}\in\mathrm{S}} P_{\mathrm{E},\mathrm{P}(g)}\, A_{\mathrm{E}}$, where $P_{\mathrm{E},\mathrm{P}(g)} = \langle C_{\mathrm{E},\mathrm{P}(g)} \rangle$ is the contact probability between enhancer E and promoter $\mathrm{P}(g)$.

In the current formulation, the ensemble-averaged AMC score is therefore equivalent to the ABC score. The added value of the AMC formulation is that it provides a conformation-level score, enabling analysis of the distribution of AMC score that explicitly accounts for multiway contact contributions.

Other definitions of the AMC score are possible. In particular, one could explicitly weight the multiway contacts beyond a simple sum of pairwise contributions. For instance, a formulation that assigns higher weight to conformations in which multiple enhancers contact the promoter simultaneously (cooperativity effects) could capture cooperative effects that are not captured by pairwise contact probabilities alone. Such alternative formulations would yield ensemble averages that differ from the ABC score and could be explored in future work to test whether explicit multiway weighting improves the correlation with gene expression.

**Breast cancer progression**

We analyzed the alterations in the chromatin organization to assess the impact on gene expression changes the MCF10 breast cancer progression consisting of cell lines MCF10A, MCF10AT1, and MCF10CA1a [20]. The analysis focused on five genes, *SPRY1*, *SCNN1G*, *SCNN1B*, *COL12A1*, and *WNT5A*, which are highlighted in [20]. To assess the relevance of the multiway enhancer-promoter contacts, we executed the following steps. (1) After preprocessing the Micro-C data, we generated the ensemble of 3D structures using the HIPPS method. (2) We determined the location of the putative enhancers. (3) We computed the activity associated with each enhancer. (4) The 3D structures allowed us to calculate the multiway enhancer-promoter contacts and the AMC score (Eq. 13). The summary statistics for all enhancer candidates are provided in Supplementary File Table 1.

**Preprocessing the Micro-C data**: We used the Micro-C contact maps for the three cell lines in MCF10 progression from the GEO accession GSE320319. For each cell line, eight replicates of Micro-C contact maps are used to construct the contact map for the selected genomic region. The raw contact matrices are extracted at 5-kb resolution using the observed KR-normalized contacts. The contact maps from the eight replicates are summed to obtain a single experimental contact map for each region and cell line. The resulting maps are coarse-grained to 10-kb resolution, which defines the locus size used in subsequent HIPPS applications. Thus, all the promoter bins, enhancer bins, and HIPPS structures are based on a common 10-kb resolution.

**Enhancer locations**: Enhancer bins were determined from H3K27ac broadPeak data from GEO accession GSE229295. For a given genomic region, the broadPeak intervals were projected onto the same 10-kb binning used for the contact maps. A bin is then associated with an enhancer if it overlaps with at least one H3K27ac broadPeak interval and does not overlap the gene body at interest. In this way, the genomic position of each enhancer is represented by the corresponding 10-kb bin.

**Enhancer activity**: For each enhancer bin, we assigned an activity using the `signalValue` field in the H3K27ac broadPeak file from GEO accession GSE229295. For a broadPeak interval overlapping a 10-kb bin, the contribution of this peak to the bin is computed as,

$$\texttt{signalValue} \times \frac{\texttt{overlap}_{bp}}{\texttt{bin length}_{bp}}, \tag{15}$$

where $\texttt{overlap}_{bp}$ is the overlap length between the broadPeak interval and the bin, and $\texttt{bin length}_{bp}$ = 10,000. The activity of the enhancer bin is then computed by summing the contributions from all broadPeak intervals that overlap with this bin.

**Enhancer-promoter contacts**: For each gene, the promoter position was determined from the transcription start site (TSS). The promoter was identified as the 10-kb bin containing the TSS. Using the HIPPS-derived connectivity matrix, we generated an ensemble of 10,000 conformations for each cell line. The contacts between the promoter and enhancers are computed using a cutoff distance $r_c = 0.2$. The average E-P 3D distances across all three cell lines for the five genes is 0.35. The average standard deviation of distances is 0.15. The cut off depends on the resolution of contact map, which is higher in the Micro-C compared to the standard Hi-C used to obtain data for prostate cancer cells [9]. Unlike in the case of prostate cancer, we could not determine the physical unit $r_c$. We varied $r_c$ in the range of 0.05 and 0.35. Fig. SI-24 shows that changing $r_c$ values in this range does not alter the conclusion, namely, that the gene expression is positively correlated with mean AMC score except for *WNT5A* gene.

**AMC score**: Following the procedure used in the context of prostate cancer (described in section **Activity-by-multiway-contact (AMC) model**), we calculated AMC score for a given conformation,

$$\text{AMC} = \sum_{\text{E}\in\text{S}} A_{\text{E}} C_{\text{E},\text{P}(g)}, \tag{16}$$

where the sum over E runs over the set of enhancers S associated with the gene $g$. The activity of enhancer E is $A_{\text{E}}$, and $C_{\text{E},\text{P}(g)}$ is the indicator function of contact between enhancer E and promoter P$(g)$. Here, $C_{\text{E},\text{P}(g)} = 1$ if the enhancer-promoter distance is smaller than the cutoff distance $r_c = 0.2$, and $C_{\text{E},\text{P}(g)} = 0$ otherwise. For each cell line, the mean AMC score is computed by averaging over the ensemble of 10,000 conformations.

TABLE SI-1: Percentage of structures in cluster-1 and cluster-2 for the 4 cell lines.

| | RWPE1 | MDAPCa2b | C42B | 22Rv1 |
|---|---|---|---|---|
| cluster-1 | 84 | 68 | 81 | 58 |
| cluster-2 | 17 | 32 | 19 | 48 |

TABLE SI-2: Gemonic position of AR promoter and enhancers in ChrX. Taken from supplementary data-5 of Rhie et al. [9].

| | Genomic position | genomic distance from Promoter |
|---|---|---|
| P | 66.76 Mb | 0 Mb |
| E1 | 66.64 Mb | 0.12 Mb |
| E2 | 66.32 Mb | 0.44 Mb |
| E3 | 66.24 Mb | 0.52 Mb |
| E4 | 66.12 Mb | 0.64 Mb |
| E5 | 66.08 Mb | 0.68 Mb |
| E6 | 66.00 Mb | 0.76 Mb |
| E7 | 65.96 Mb | 0.80 Mb |
| E8 | 65.92 Mb | 0.84 Mb |

TABLE SI-3: Gemonic position of FOXA1 promoter and enhancers in Chr14. Taken from supplementary data-5 of Rhie et al. [9].

| | Genomic position | genomic distance from Promoter |
|---|---|---|
| P | 38.04 Mb | 0 Mb |
| E1 | 37.88 Mb | -0.16 Mb |
| E2 | 37.76 Mb | -0.28 Mb |
| E3 | 38.32 Mb | 0.28 Mb |
| E4 | 37.72 Mb | -0.32 Mb |
| E5 | 37.68 Mb | -0.36 Mb |
| E6 | 39.44 Mb | 1.40 Mb |
| E7 | 39.48 Mb | 1.44 Mb |
| E8 | 39.52 Mb | 1.48 Mb |

TABLE SI-4: Epigenetic details of the common TADs (Topologically Associating Domains) sharing boundaries in the normal RWPE1 and cancerous 22Rv1 cell lines. Number of active (A) and repressed (B) TADs for each chromosome from 1 to X. List of the number of TADs that change their epigenetic state from B to A and vice versa, as well as the number of TADs that maintain their epigenetic state.

| Chr no. | RWPE1 | | 22Rv1 | | B to A | A to B | A to A | B to B |
|---|---|---|---|---|---|---|---|---|
| | A type | B type | A type | B type | | | | |
| 1 | 49 | 194 | 50 | 193 | 9 | 8 | 41 | 185 |
| 2 | 29 | 165 | 22 | 172 | 4 | 11 | 18 | 161 |
| 3 | 19 | 149 | 32 | 136 | 17 | 4 | 15 | 132 |
| 4 | 14 | 154 | 9 | 159 | 2 | 7 | 7 | 152 |
| 5 | 23 | 160 | 22 | 161 | 5 | 6 | 17 | 155 |
| 6 | 21 | 154 | 18 | 157 | 5 | 8 | 13 | 149 |
| 7 | 6 | 136 | 21 | 121 | 16 | 1 | 5 | 120 |
| 8 | 8 | 127 | 13 | 122 | 7 | 2 | 6 | 120 |
| 9 | 11 | 99 | 22 | 88 | 11 | 0 | 11 | 88 |
| 10 | 24 | 79 | 17 | 86 | 0 | 7 | 17 | 79 |
| 11 | 22 | 113 | 24 | 111 | 6 | 4 | 18 | 107 |
| 12 | 24 | 121 | 30 | 115 | 11 | 5 | 19 | 110 |
| 13 | 8 | 63 | 6 | 65 | 0 | 2 | 6 | 63 |
| 14 | 20 | 53 | 19 | 54 | 1 | 2 | 18 | 52 |
| 15 | 11 | 60 | 20 | 51 | 9 | 0 | 11 | 51 |
| 16 | 21 | 52 | 21 | 52 | 2 | 2 | 19 | 50 |
| 17 | 23 | 63 | 30 | 56 | 12 | 5 | 18 | 51 |
| 18 | 7 | 68 | 10 | 65 | 5 | 2 | 5 | 63 |
| 19 | 13 | 31 | 22 | 22 | 9 | 0 | 13 | 22 |
| 20 | 14 | 56 | 17 | 53 | 5 | 2 | 12 | 51 |
| 21 | 7 | 36 | 8 | 35 | 2 | 1 | 6 | 34 |
| 22 | 6 | 33 | 22 | 17 | 16 | 0 | 6 | 17 |
| X | 1 | 110 | 0 | 111 | 0 | 1 | 0 | 110 |

TABLE SI-5: Epigenetics of the common TADs (Topologically Associating Domains) sharing boundaries in the normal RWPE1 and cancerous C42B cell lines. List of the number of active (A) and repressed (B) TADs for each chromosome from 1 to X. It also details the number of TADs that change their epigenetic state from B to A and vice versa, as well as the number of TADs that maintain their epigenetic state.

| Chr no. | RWPE1 | | C42B | | B to A | A to B | A to A | B to B |
|---|---|---|---|---|---|---|---|---|
| | A type | B type | A type | B type | | | | |
| 1 | 49 | 182 | 46 | 185 | 12 | 15 | 34 | 170 |
| 2 | 27 | 181 | 27 | 181 | 12 | 12 | 15 | 169 |
| 3 | 21 | 151 | 32 | 140 | 15 | 4 | 17 | 136 |
| 4 | 14 | 136 | 18 | 132 | 9 | 5 | 9 | 127 |
| 5 | 19 | 151 | 22 | 148 | 9 | 6 | 13 | 142 |
| 6 | 18 | 157 | 13 | 162 | 3 | 8 | 10 | 154 |
| 7 | 8 | 132 | 20 | 120 | 13 | 1 | 7 | 119 |
| 8 | 10 | 114 | 17 | 107 | 9 | 2 | 8 | 105 |
| 9 | 11 | 110 | 18 | 103 | 10 | 3 | 8 | 100 |
| 10 | 19 | 97 | 19 | 97 | 6 | 6 | 13 | 91 |
| 11 | 21 | 121 | 22 | 120 | 6 | 5 | 16 | 115 |
| 12 | 28 | 107 | 28 | 107 | 6 | 6 | 22 | 101 |
| 13 | 8 | 51 | 6 | 53 | 4 | 6 | 2 | 47 |
| 14 | 17 | 50 | 18 | 49 | 5 | 4 | 13 | 45 |
| 15 | 14 | 73 | 20 | 67 | 8 | 2 | 12 | 65 |
| 16 | 22 | 55 | 22 | 55 | 5 | 5 | 17 | 50 |
| 17 | 22 | 63 | 22 | 63 | 5 | 5 | 17 | 58 |
| 18 | 5 | 57 | 11 | 51 | 6 | 0 | 5 | 51 |
| 19 | 12 | 23 | 15 | 20 | 4 | 1 | 11 | 19 |
| 20 | 19 | 53 | 19 | 53 | 3 | 3 | 16 | 50 |
| 21 | 10 | 39 | 11 | 38 | 3 | 2 | 8 | 36 |
| 22 | 7 | 31 | 17 | 21 | 11 | 1 | 6 | 20 |
| X | 3 | 109 | 12 | 100 | 10 | 1 | 2 | 99 |

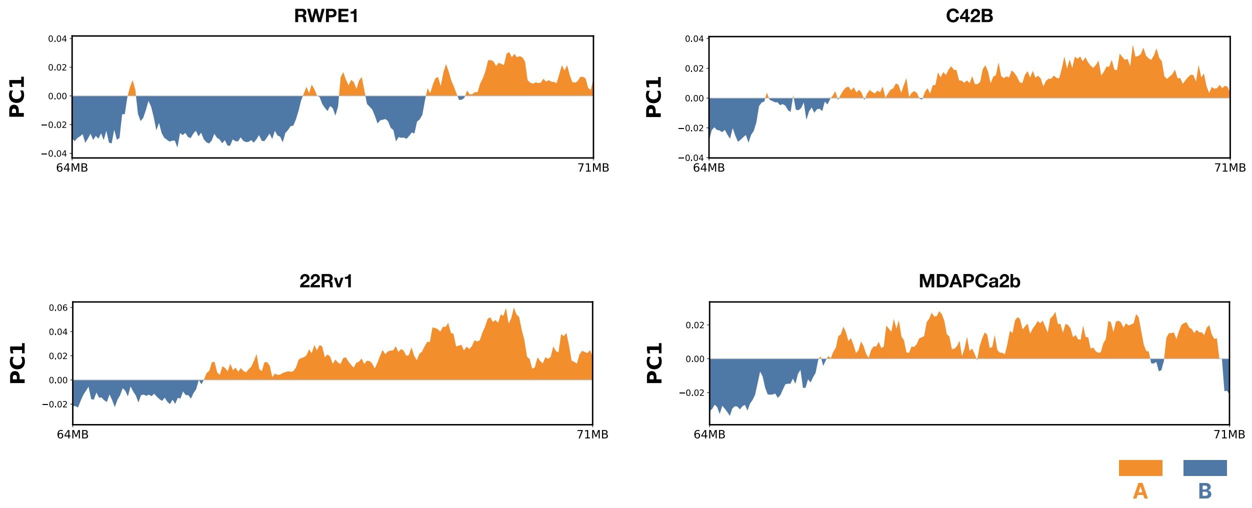


FIG. SI-1: **Assignment of epigenetic states:** Principal component 1 (PC1) profiles across chromosome X (64–71 Mb) in RWPE1, C42B, 22Rv1, and MDAPCa2b cell lines. Positive PC1 values in orange are associated with euchromatin (compartment A) with active histone marks. Negative PC1 values in blue correspond to heterochromatin (compartment B) with inactive histone marks.

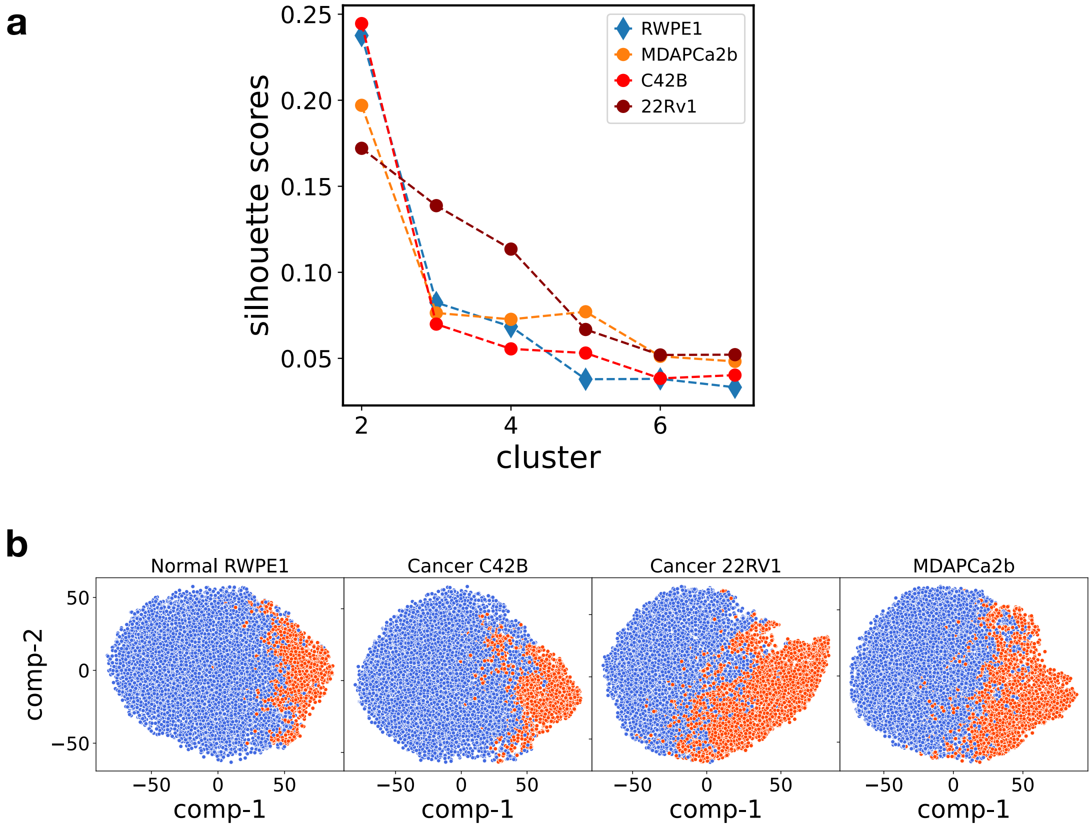


FIG. SI-2: **Clustering of the conformational ensemble:** (a) Silhouette scores for clustering in normal and cancer cell lines. Higher scores are found when the ensembles are divided into two clusters across all the cell lines. (b) Two-dimensional t-SNE representation of the structural ensembles, with blue representing cluster 1 and red representing cluster 2.

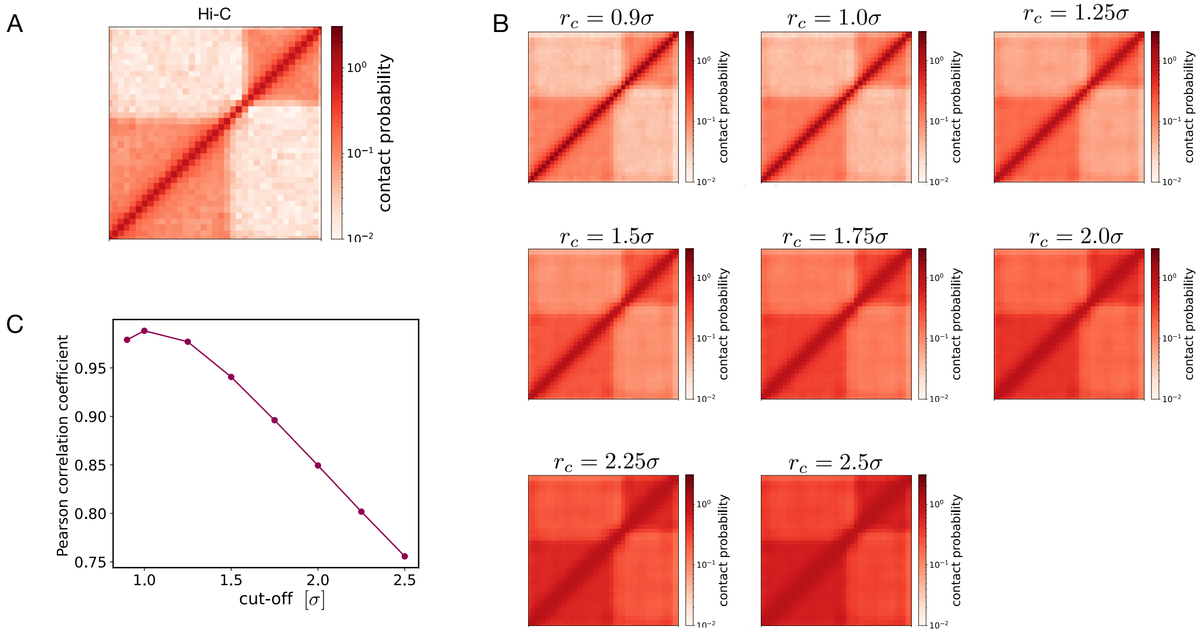


FIG. SI-3: **Threshold distance for contact:** (A) Hi-C contact map for the AR locus region. (B) Contact maps generated from the HIPPS-generated structures as a function of the contact threshold from $r_c = 0.9\sigma$ to $r_c = 2.5\sigma$. (C) Pearson correlation between the Hi-C data and HIPPS-generated contact maps as a function of $r_c$.

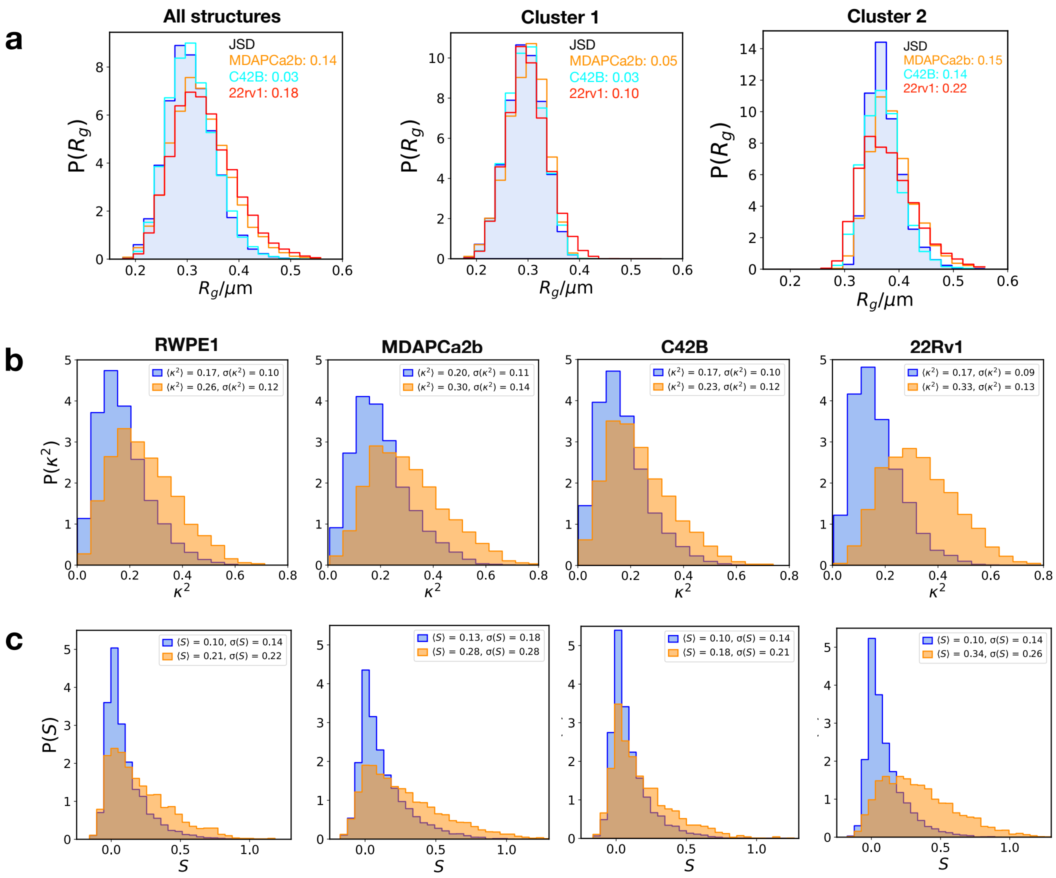

FIG. SI-4: **Structural features of chromosome X:** (a) Probability distribution of the radius of gyration $P(R_g)$. JSD, defined in Eq. 6, between the distribution of cancer cell line compared to the normal RWPE1. (b) Histogram of the shape anisotropy ($\kappa^2$) separately for cluster-1 and cluster-2 for RWPE1, MDAPCa2b, C42B and 22Rv1 cell lines. (c) Historagram of shape parameter, $S$, separately for cluster-1 and cluster-2 fore RWPE1, MDAPCa2b, C42B and 22Rv1 cell lines.

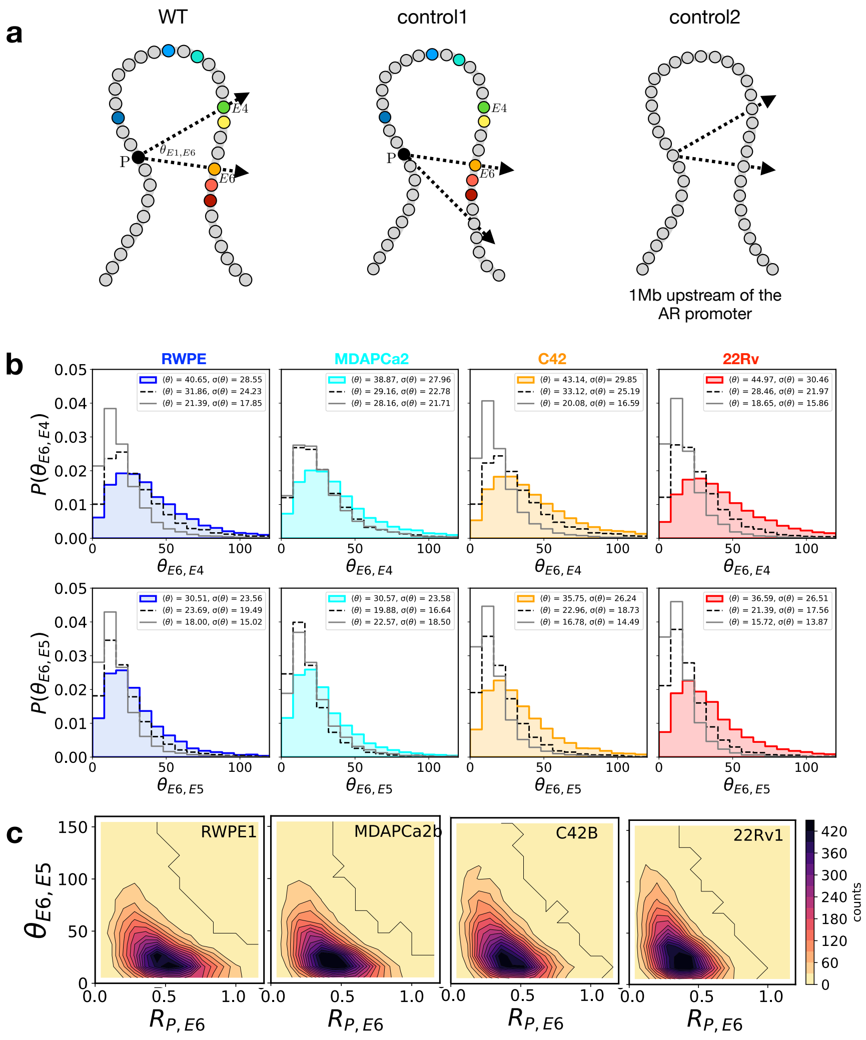


FIG. SI-5: **Angles between promoter and enhancers across cell lines do not change significantly:** (a) Schematic representation of the AR region in chromosome X, depicting the angle ($\theta_{E6,E4}$) between the promoter E6 and E4 keeping the promoter P as an anchor. The black bead denotes the AR promoter, while the enhancers are represented by colored beads ranging from blue (E1) to red (E8). The angle $\theta_{E6,E4}$ is formed between the vector from the promoter (P) to enhancer E6 and the vector from the promoter (P) to enhancer E4. control 1: the angle between E6 and a locus that is at the same genomic distance as E4 from E6, keeping P as the anchor. control 2: for control 2, we selected a triplet of loci situated 1 Mb upstream of the AR promoter. This represents a scenario where none of the loci act as enhancers or the promoter for AR, while maintaining the genomic distance among the loci. (b) Top panel: Probability distribution of the angle $\theta_{E6,E4}$ across different cell lines. The black dashed (gray solid) line is control 1 (control 2). (c) 2D contour plot between the distance $R_{P,E6}$ and angle $\theta_{E6,E5}$.

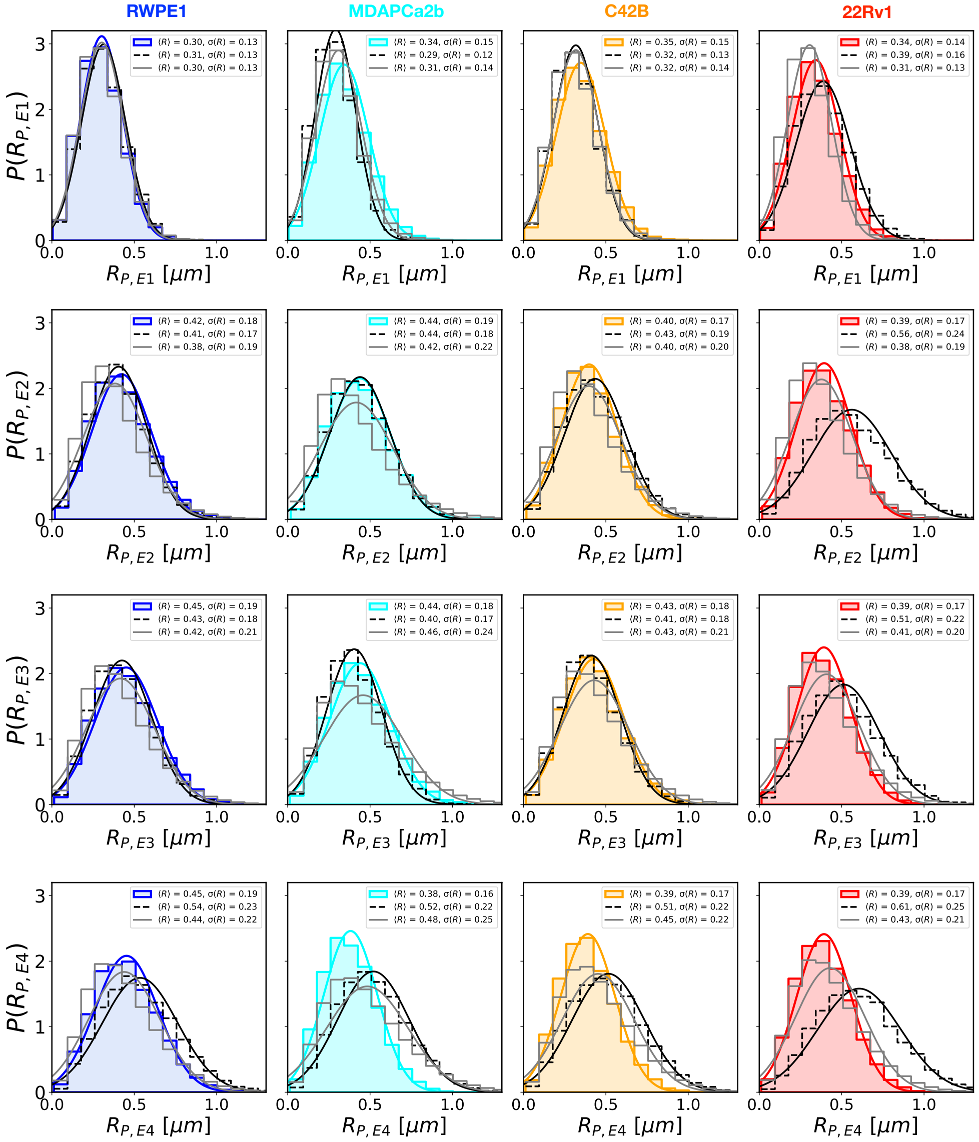


FIG. SI-6: **Distance distributions across cell lines for AR locus:** Each panel shows probability distributions of E-P distances for the AR gene for enhancers E1 to E4 with the promoter in the four cell lines labeled on top. Black lines correspond to promoter-locus distance distribution at the same genomic distance as a given enhancer. Grey lines represent distance distributions for two randomly chosen loci whose genomic distance matches that between the promoter and a given enhancer. The mean values and the standard deviations are given the panels.

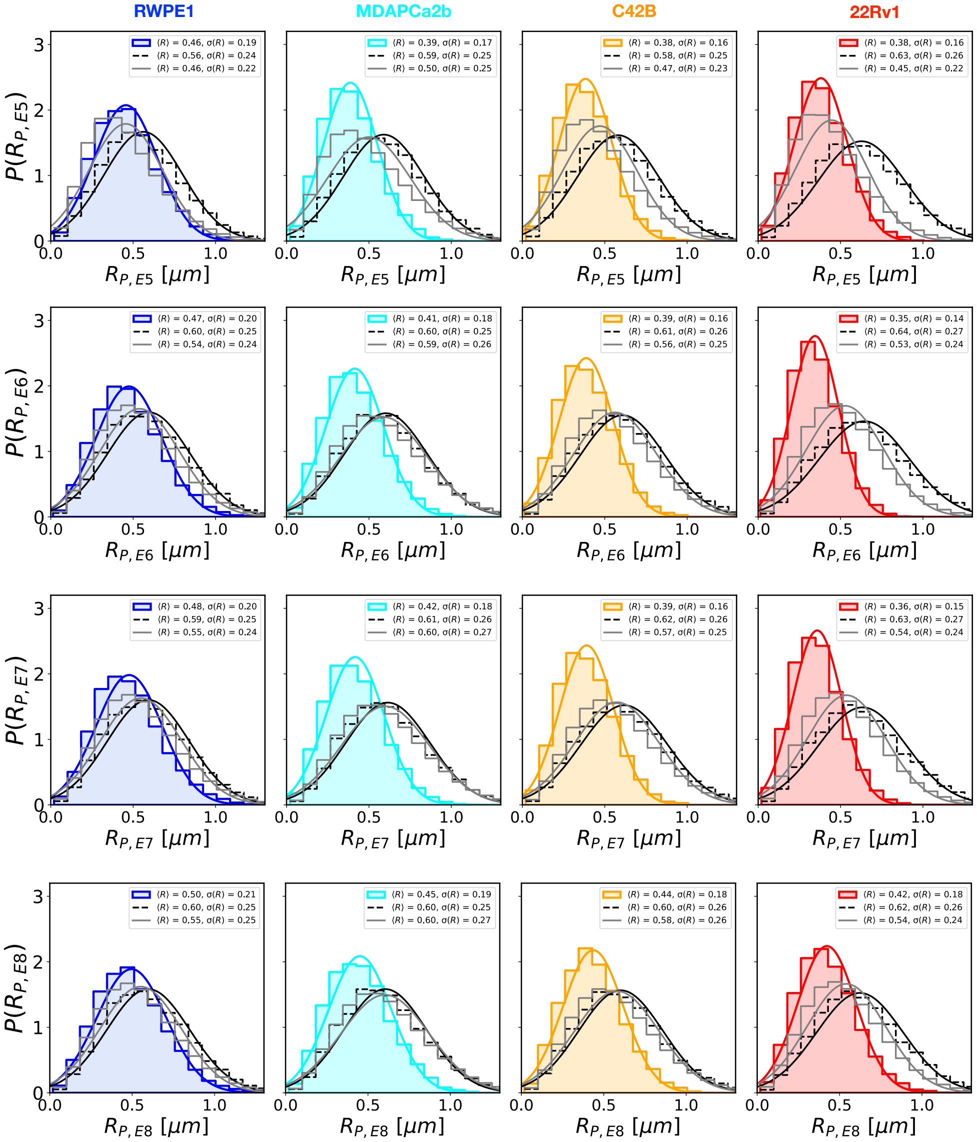


FIG. SI-7: **Distance distributions between Enhancers and promoter in AR locus:** Same as Fig. SI-6 except these correspond to E-P distance of AR gene for enhancer E5 to E8 with the promoter.

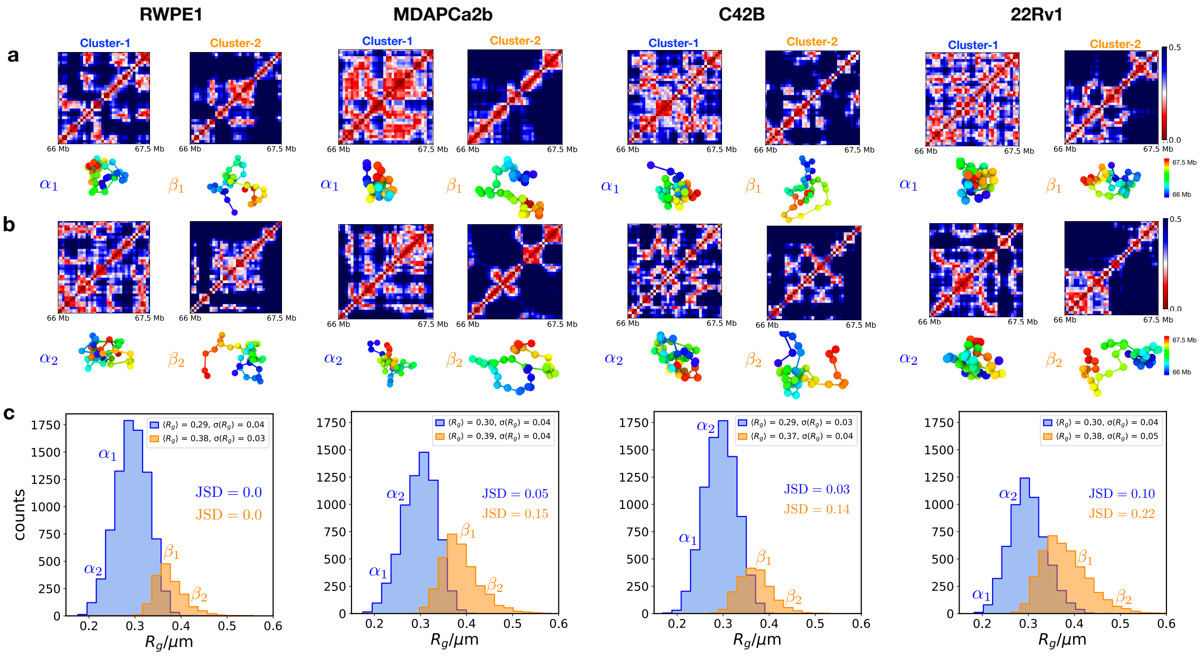


FIG. SI-8: **Structural heterogeneity in the AR locus region varies with the cell type:** (a) Distance maps for the HIPPS-inferred chromatin conformations corresponding to two distinct groups, cluster-1 (left) and cluster-2 (right), which are determined using the clustering algorithm described in the SI. The corresponding chromatin structures for each distance map are also shown. Distance maps and structures are presented for all four cell lines studied. (b) Similar to panel a, this panel shows an alternative distance map and structure for each cluster separately. (c) Probability distribution of the radius of gyration ($R_g$) of the chromatin conformations, compared between cluster-1 (blue) and cluster-2 (orange) across different cell lines. $\alpha$ and $\beta$ denote the $R_g$ values of the structures depicted in panels a and b.

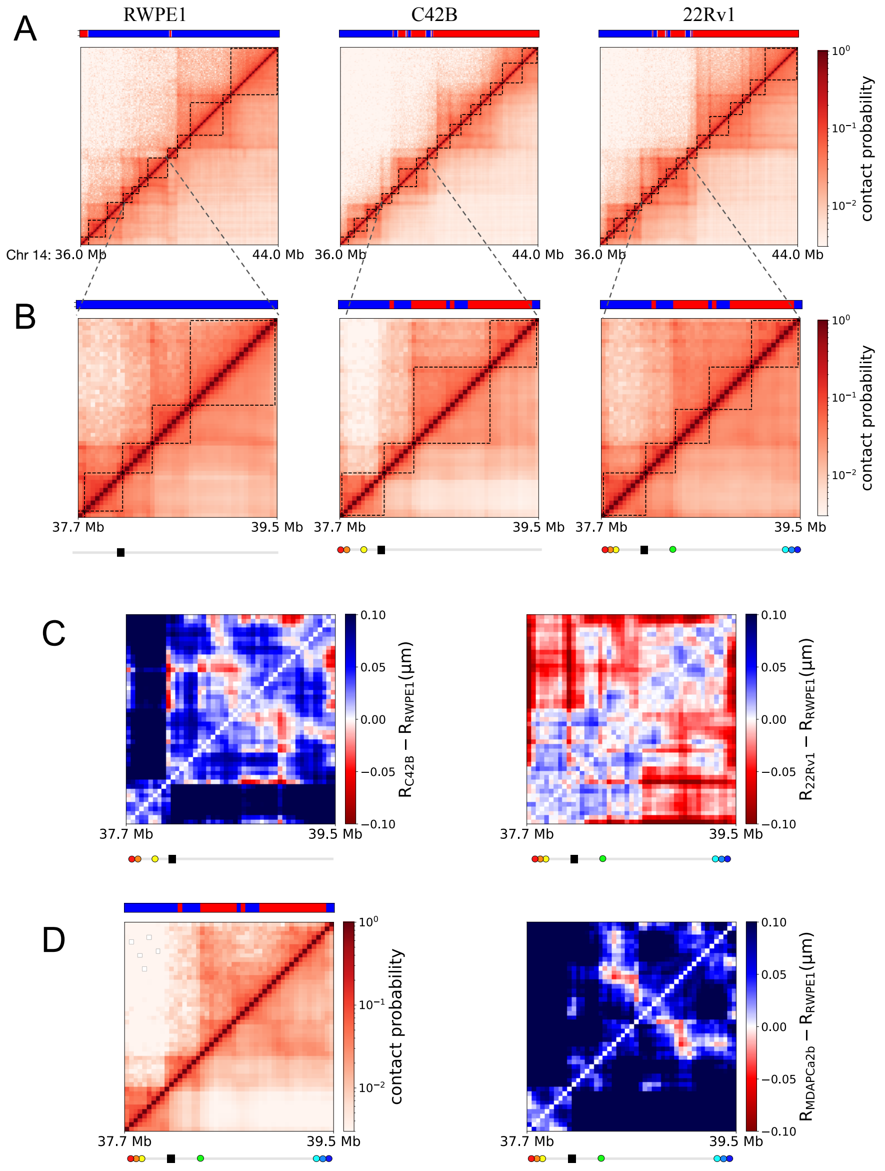


FIG. SI-9: **FOXA1:** (A) Comparison of the calculated contact map using HIPPS and Hi-C for a 8MB region in Chr14 for the normal RWPE1 and cancer cell (C42B and 22Rv1) lines. (B) Zoomed region of in Chr14 containing FOXA1 promoter and enhancers. The location of enhancers and the promoter is shown by the black rectangle and the colored circle below. (C) Distance map difference between the cancer cell line C42B and 22Rv1 and the normal RWPE1. (D) Contact map comparison with the Hi-C and distance map difference with RWPE1 for cancer cell line MDAPCa2b.

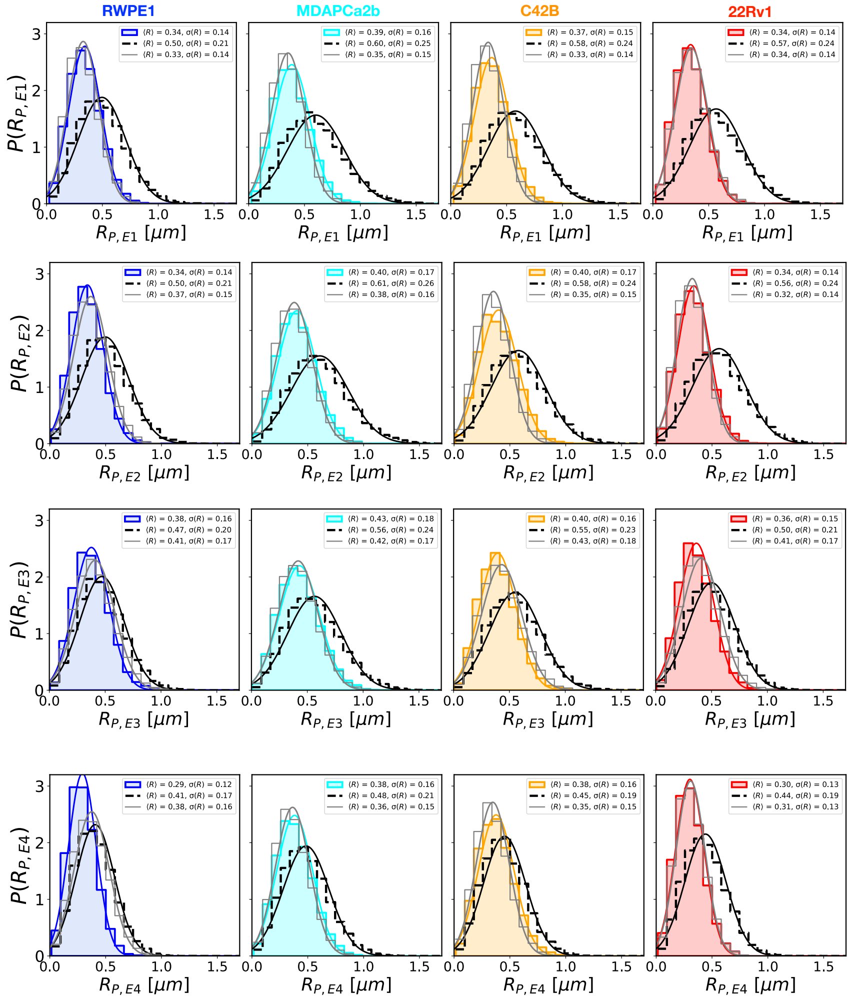


FIG. SI-10: **Distance distributions involving FOXA1:** Distribution of E-P distances for enhancers E1 to E4 with the FOXA1 promoter. Black lines (control 1) correspond to promoter-locus distance distribution at the same genomic distance as a given enhancer. Grey lines (control 2) represent distance distributions for two randomly chosen loci whose genomic distance matches that between the promoter and a given enhancer. The mean values and the standard deviations are given the panels.

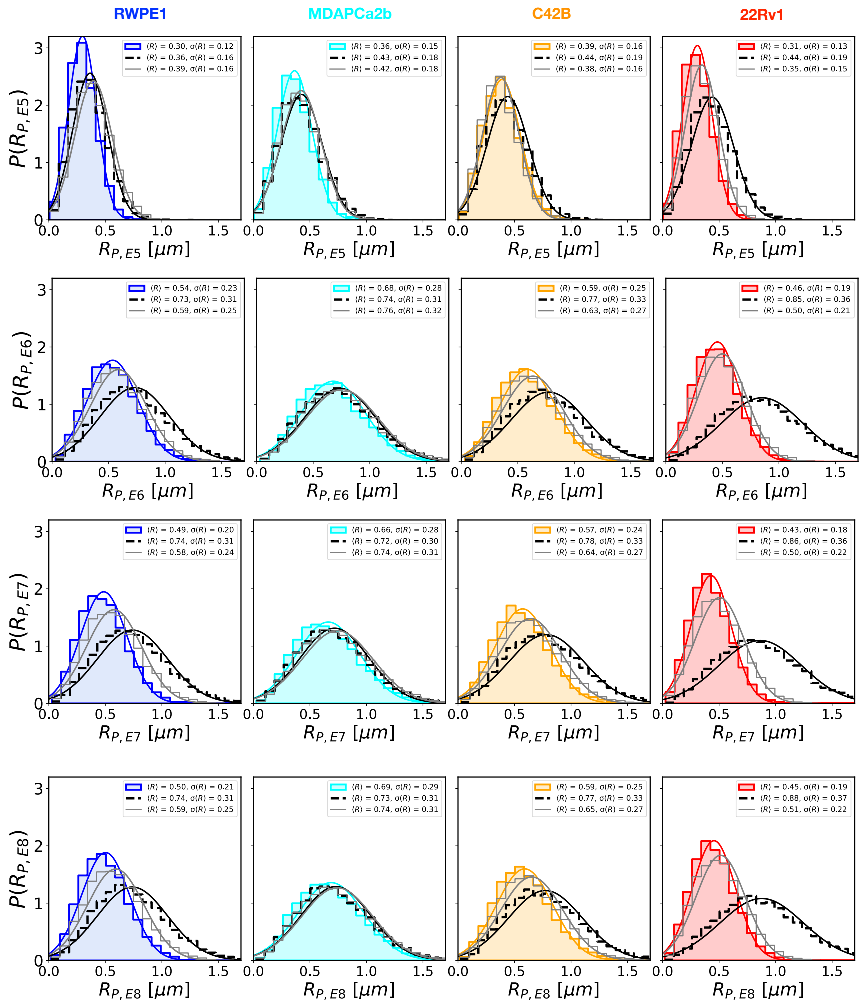


FIG. SI-11: **Distance distributions for FOXA1:** Same as Fig. SI-10 except these involve distances between the promoter and enhancers E5 to E8.

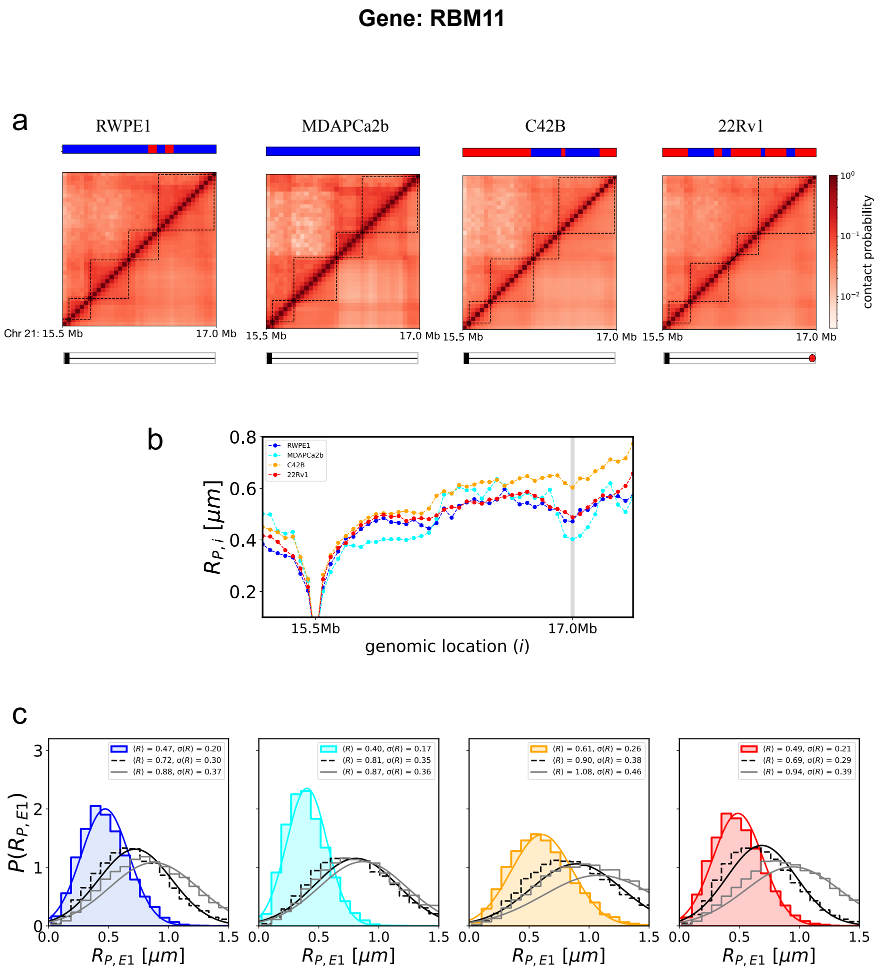


FIG. SI-12: **Gene RBM11**: (a) Comparison of contact maps from HIPPS and Hi-C for a 1.5 MB region on chromosome 21, highlighting the normal RWPE1 cell line and cancer cell lines C42B, 22Rv1, and MDAPCa2b. (b) Average distance, in $\mu$m, from the AR promoter to the $i^{th}$ locus in the chromosome 21 region spanning 15.3 Mb to 17.2 Mb, with genomic distance measured from the promoter. The gray vertical lines indicate the locations of enhancers. (c) Distance distribution between the RBM11 gene promoter (P) and the sole enhancer E1 across different cell lines, represented in various colors. The black dashed line denotes Control 1, reflecting the distance distribution between P and a locus at the same genomic distance as E1. The gray solid line represents Control 2, corresponding to two arbitrary loci with the same genomic distance as that between the promoter and E1. The averages were calculated over all such pairs.

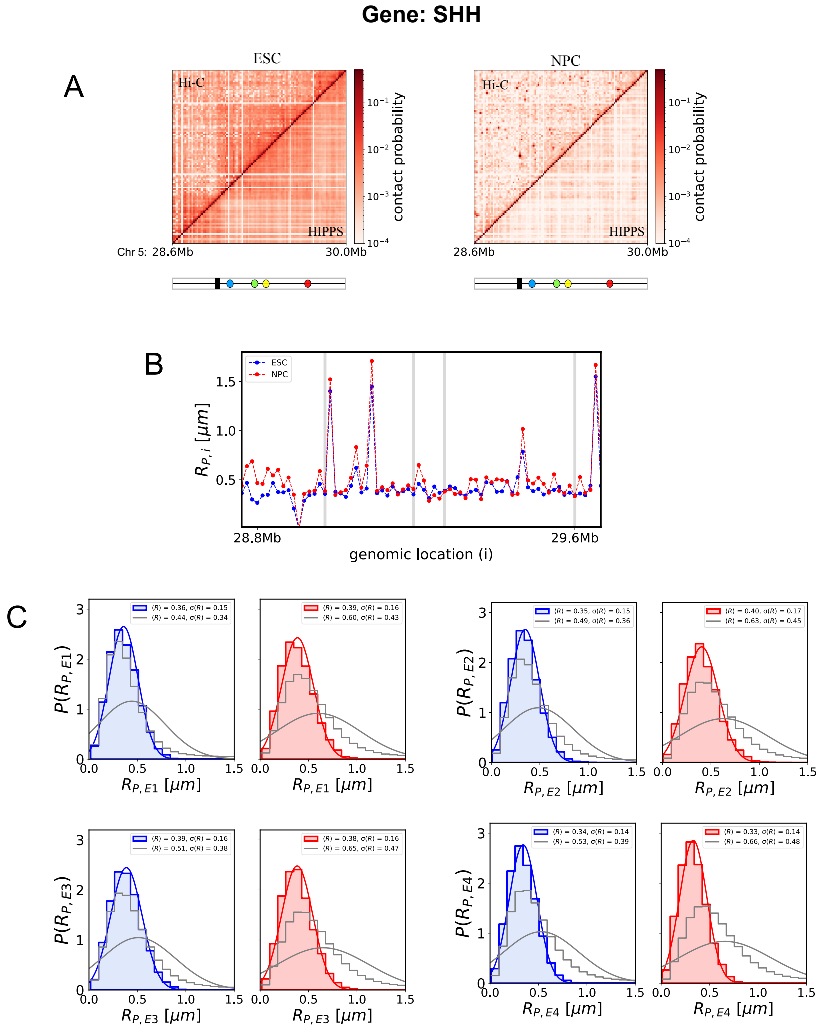


FIG. SI-13: **SHH gene locus:** (A) Comparison of contact maps from HIPPS and 5C for a 1.6 MB region on chromosome 5 for the ESC and NPC cell lines. (B) Average distance, in $\mu$m, from the SHH promoter to the $i^{th}$ locus in the chromosome 5 region spanning 28.6 Mb to 30.0 Mb, with genomic distance measured from the promoter. The gray vertical lines indicate the locations of enhancers. (C) Probability distribution of distances between the SHH promoter and its four enhancers (E1 to E4). Blue and red represent the ESC and NPC cell lines, respectively. The gray solid line corresponds to Control 2, reflecting two arbitrary loci with the same genomic distance as that between the promoter and E1. Averages were calculated over all such pairs.

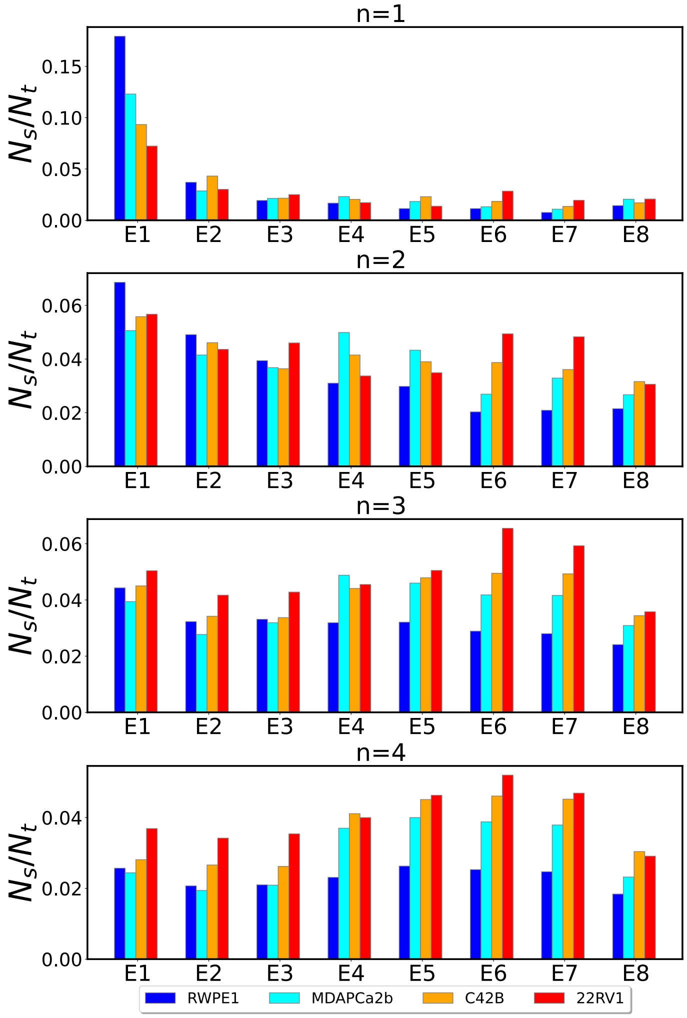


FIG. SI-14: **Enhancer contributions for the AR locus:** The contribution of each enhancer when $n = 1, 2, 3, 4$ enhancers is in contact with the promoter for the AR locus. Although multiple enhancers are in contact with the promoter their contribution decreases as the number of multiway contacts increases.

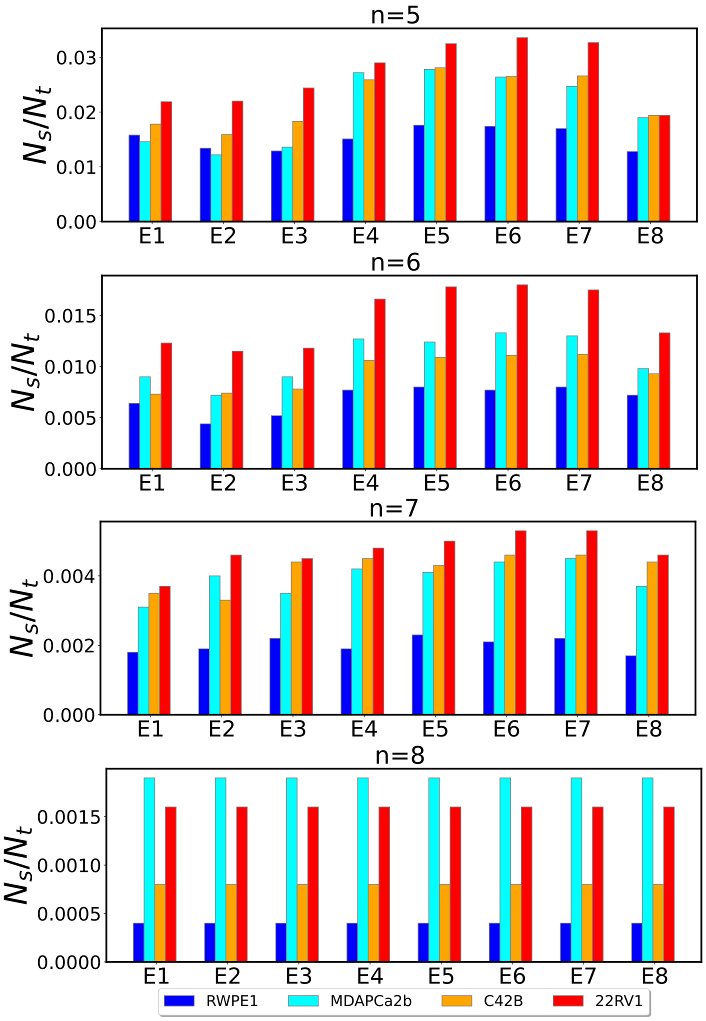


FIG. SI-15: **Multiway contributions to E-P interactions:** The contribution of each enhancer for the cases when $n = 5, 6, 7, 8$ enhancers is in contact with the promoter for the AR locus. Clearly, for $n > 4$ the contribution is negligible.

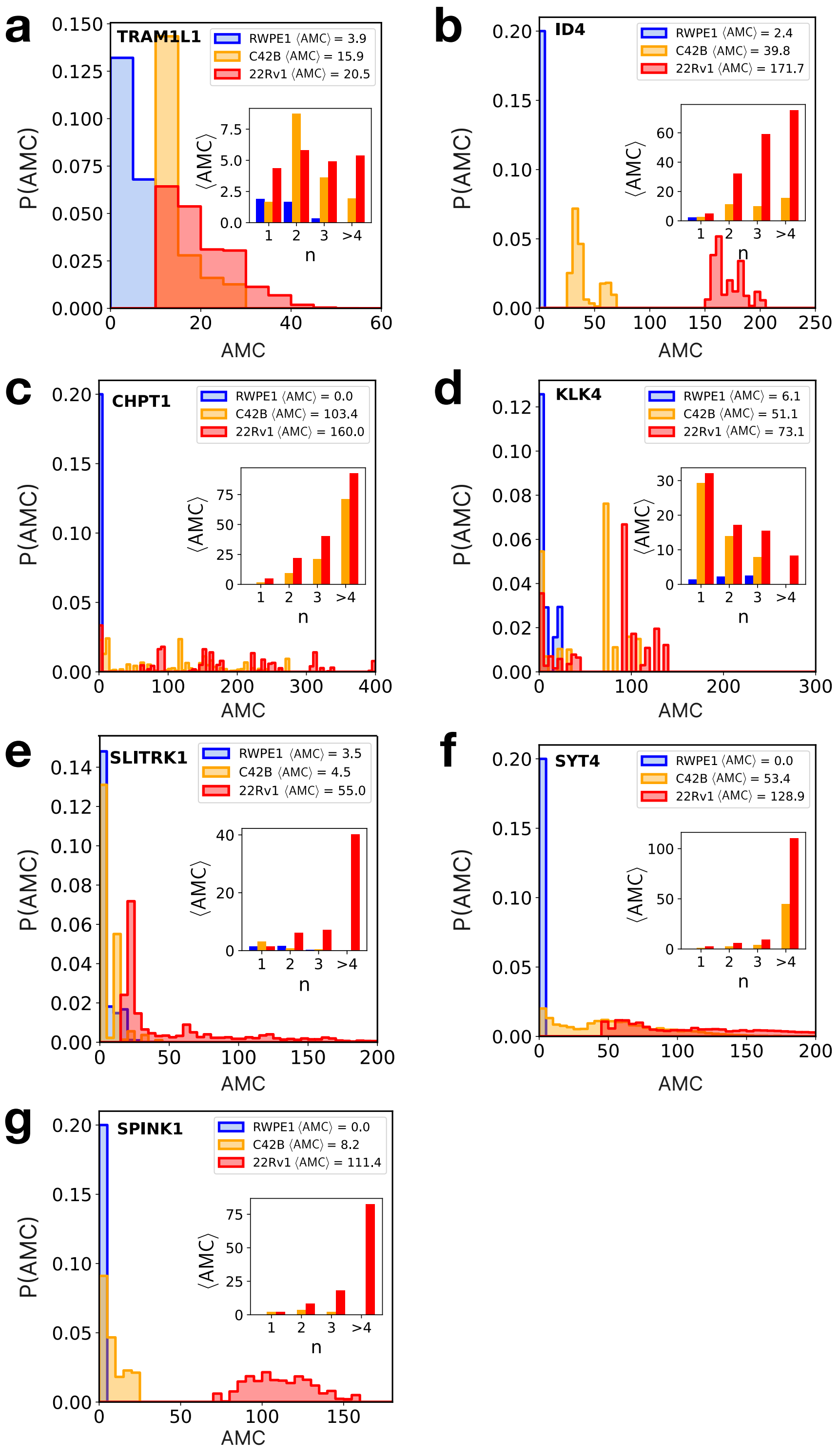


FIG. SI-16: **AMC scores:** (a-g) Distribution of AMC scores for several genes. The inset shows the contribution of multi-way contacts involving $n$ enhancers to the mean AMC score for each cell line. The AMC scores correlate well with the gene expression.

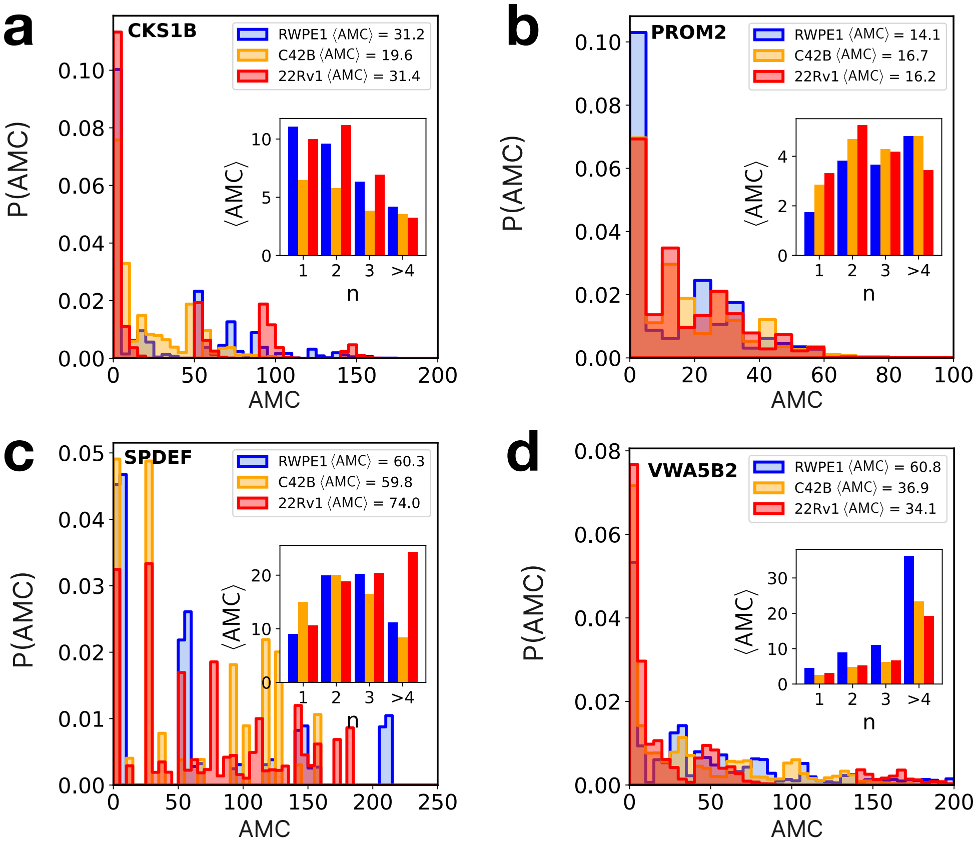


FIG. SI-17: **AMC scores:** (a-d) Distribution of AMC scores for the genes whose expression levels are not explained well by the scores. The inset shows the contribution of multi-way contacts involving $n$ enhancers to the mean AMC score for each cell line.

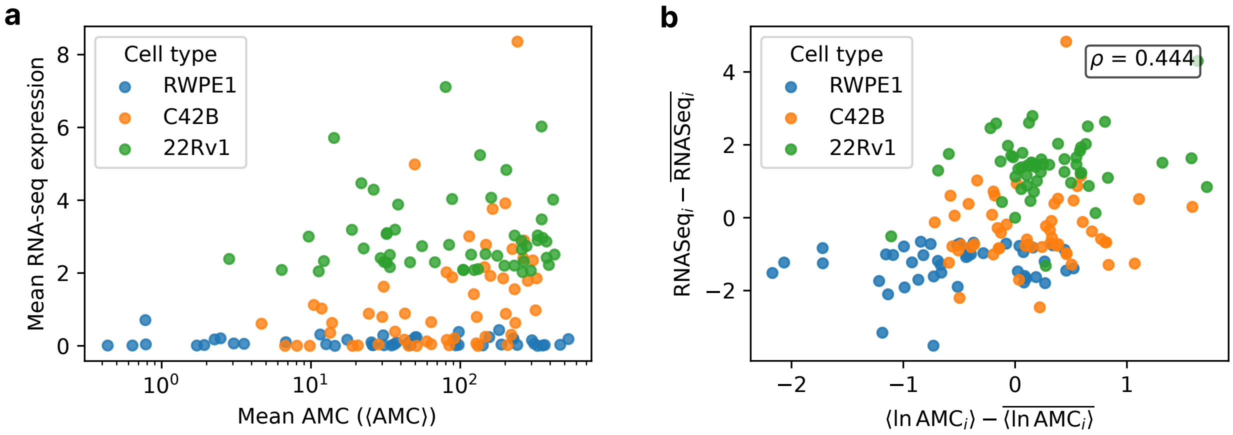


FIG. SI-18: **Correlation between AMC scores and RNA-seq expression for the 54 genes in prostate cancer with enhancer information from the full 162 gene pool:** (a) Mean RNA-seq expression versus mean AMC score ($\langle \text{AMC} \rangle$) for the 54 genes across different cell lines. Each point represents a gene, with colors indicating cell lines: blue (RWPE1), orange (C42B), and red (22Rv1). (b) Gene-centered analysis showing RNA-seq - $\overline{\text{RNA-seq}}$ versus $\langle \text{AMC} \rangle - \overline{\langle \text{AMC} \rangle}$, where the overbar represents the average over the three cell lines for each gene. The analysis removes gene-specific baseline effects to focus on cell-type-specific variations. Although not precise, the correlation coefficient $\rho \approx 0.45$ is statistically significant ($p = 8.2 \times 10^{-9}$).

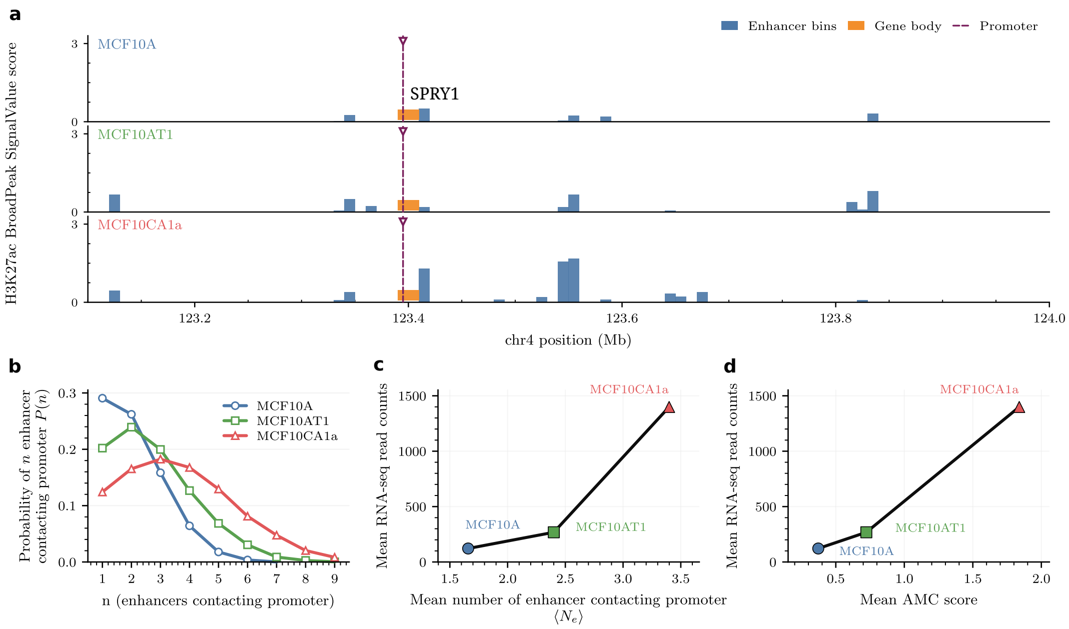


FIG. SI-19: **Enhancer-promoter contacts and enhancer activity at the *SPRY1* locus across the MCF10 breast cancer progression.** (a) Linear genomic track across the *SPRY1* locus showing the H3K27ac broadPeak `signalValue` score in MCF10A, MCF10AT1, and MCF10CA1a cell lines. Putative enhancer bins are indicated together with the gene body and promoter position. (b) Distribution of the number of enhancers contacting the promoter, $P(n)$, calculated using the HIPPS-generated structural ensemble for each cell line. (c) Mean RNA-seq read counts as a function of the average number of enhancers contacting the promoter, $\langle N_e \rangle$, for MCF10A, MCF10AT1, and MCF10CA1a. (d) Mean RNA-seq read counts as a function of the mean AMC score for the three cell lines.

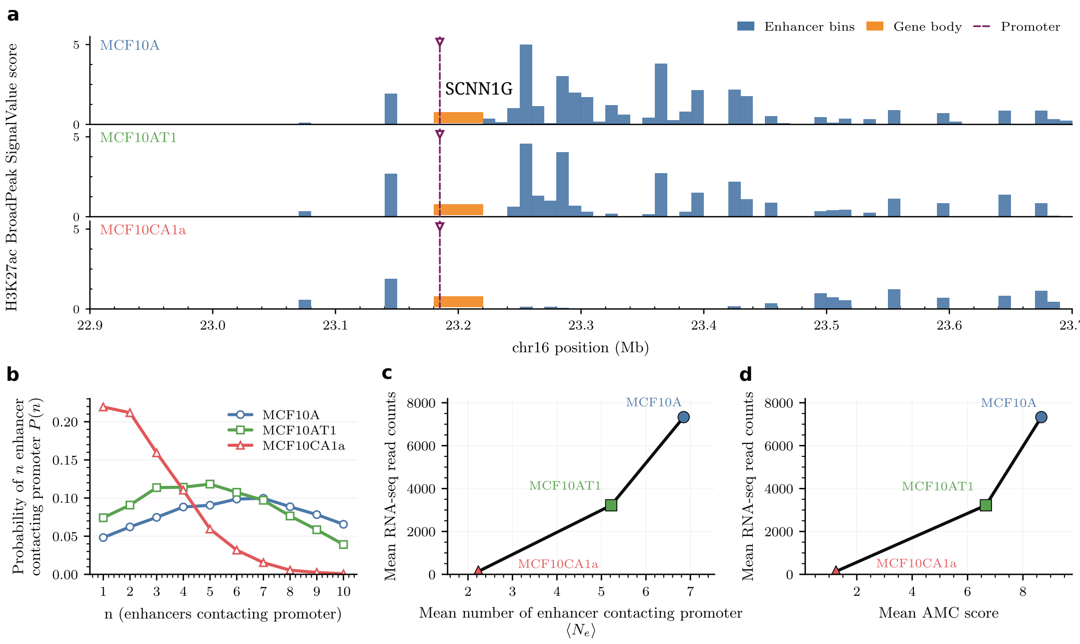


FIG. SI-20: **Enhancer-promoter contacts and enhancer activity at the *SCNN1G* locus across the MCF10 breast cancer progression.** (a) Linear genomic track across the *SCNN1G* locus showing the H3K27ac broadPeak `signalValue` score in MCF10A, MCF10AT1, and MCF10CA1a. Putative enhancer bins are indicated together with the gene body and promoter position. (b) Distribution of the number of enhancers contacting the promoter, $P(n)$, computed from the HIPPS-generated structural ensemble for each cell line. (c) Mean RNA-seq read counts as a function of the mean number of enhancers contacting the promoter, $\langle N_e \rangle$, for MCF10A, MCF10AT1, and MCF10CA1a. (d) Mean RNA-seq read counts as a function of the mean AMC score for the three cell lines.

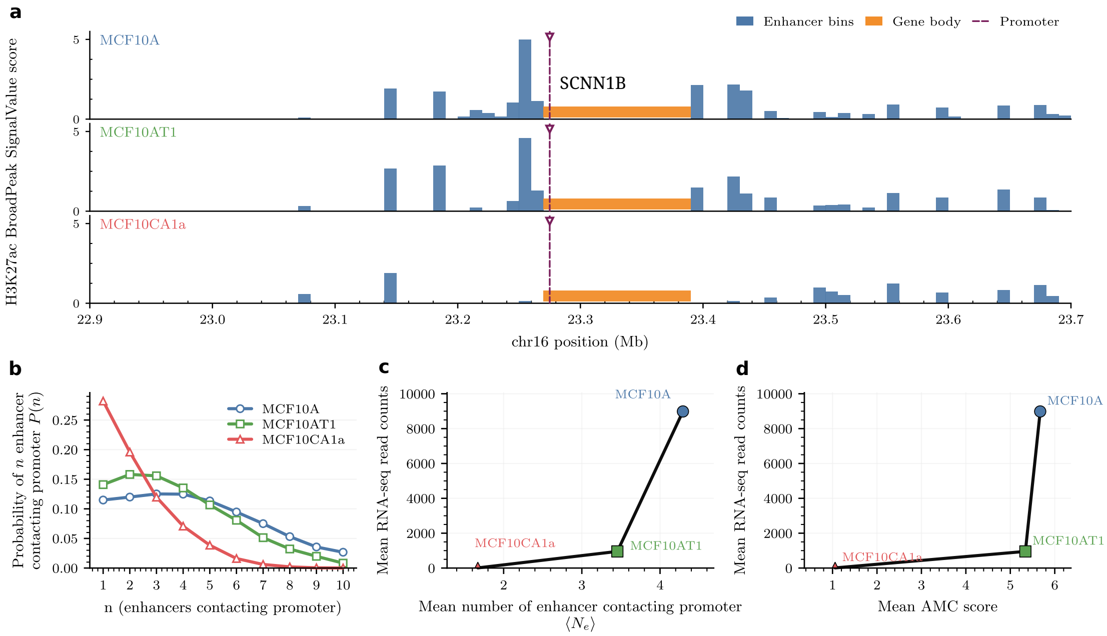


FIG. SI-21: **Enhancer-promoter contacts and enhancer activity at the *SCNN1B* locus across the MCF10 breast cancer progression.** (a) Linear genomic track across the *SCNN1B* locus showing the H3K27ac broadPeak `signalValue` score in MCF10A, MCF10AT1, and MCF10CA1a. Putative enhancer bins are indicated together with the gene body and promoter position. (b) Distribution of the number of enhancers contacting the promoter, $P(n)$, calculated using the HIPPS-generated structural ensemble for each cell line. (c) Mean RNA-seq read counts as a function of the mean number of enhancers contacting the promoter, $\langle N_e \rangle$, for MCF10A, MCF10AT1, and MCF10CA1a. (d) Mean RNA-seq read counts as a function of the mean AMC score for the three cell lines.

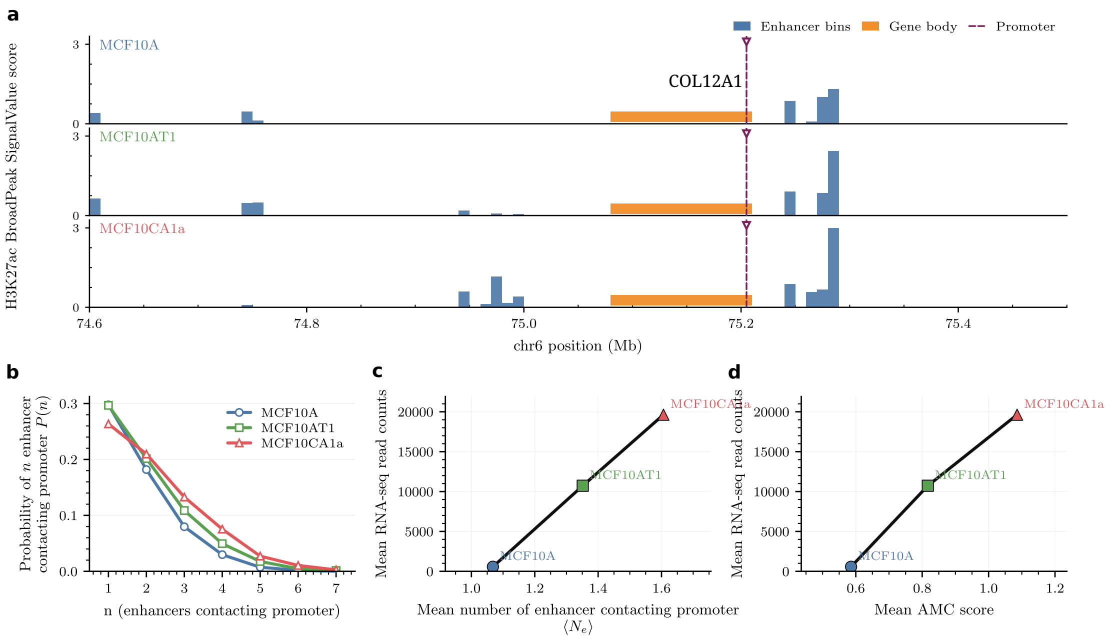


FIG. SI-22: **Enhancer-promoter contacts and enhancer activity at the *COL12A1* locus across the MCF10 breast cancer progression.** (**a**) Linear genomic track across the *COL12A1* locus showing the H3K27ac broadPeak `signalValue` score in MCF10A, MCF10AT1, and MCF10CA1a. Putative enhancer bins are indicated together with the gene body and promoter position. (b) Distribution of the number of enhancers contacting the promoter, $P(n)$, computed from the HIPPS-generated structural ensemble for each cell line. (c) Mean RNA-seq read counts as a function of the mean number of enhancers contacting the promoter, $\langle N_e \rangle$, for MCF10A, MCF10AT1, and MCF10CA1a. (d) Mean RNA-seq read counts as a function of the mean AMC score for the three cell lines.

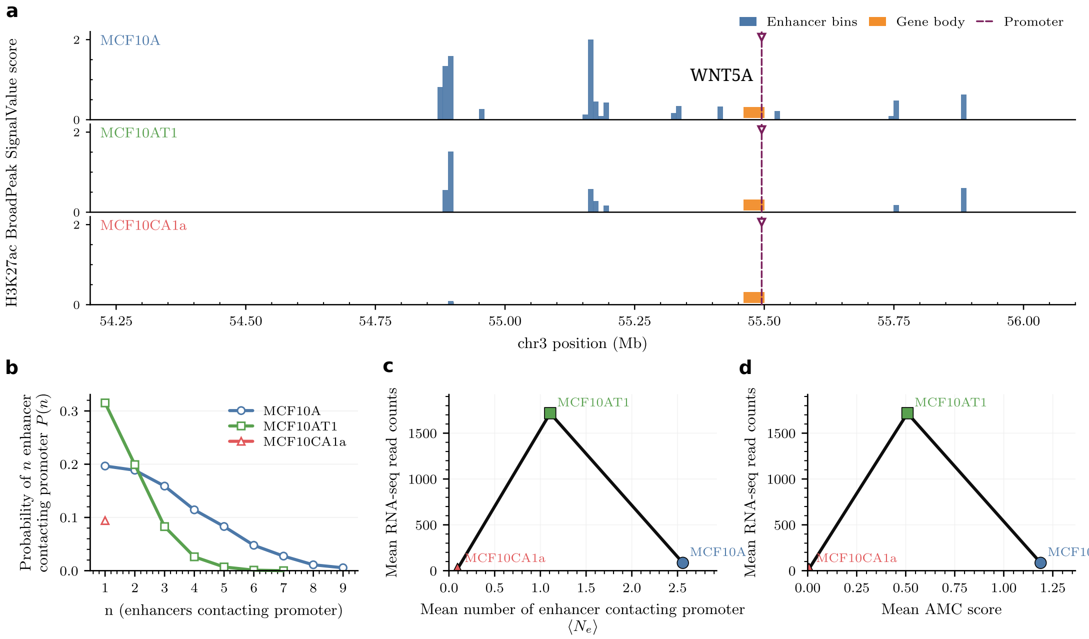


FIG. SI-23: **Enhancer-promoter contacts and enhancer activity at the *WNT5A* locus across the MCF10 breast cancer progression.** (a) Linear genomic track across the *WNT5A* locus showing the H3K27ac broadPeak `signalValue` score in MCF10A, MCF10AT1, and MCF10CA1a. Putative enhancer bins are indicated together with the gene body and promoter position. (b) Distribution of the number of enhancers contacting the promoter, $P(n)$, computed from the HIPPS-generated structural ensemble for each cell line. (c) Mean RNA-seq read counts as a function of the mean number of enhancers contacting the promoter, $\langle N_e \rangle$, for MCF10A, MCF10AT1, and MCF10CA1a. (d) Mean RNA-seq read counts as a function of the mean AMC score for the three cell lines.

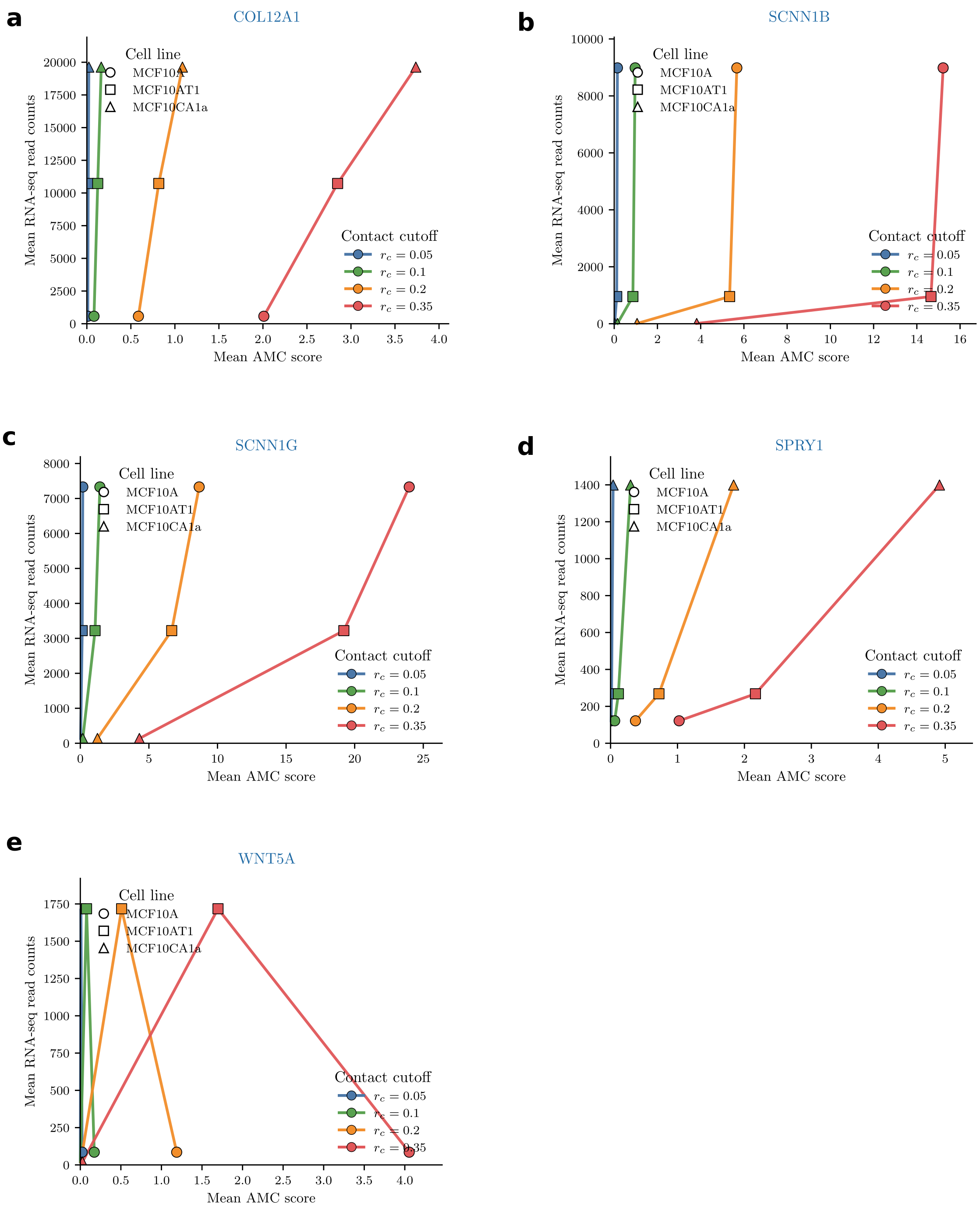


FIG. SI-24: **Effect of contact threshold $r_c$ on AMC scores.** Mean RNA-seq read counts versus the mean AMC scores computed using different values of contact threshold $r_c$ for breast cancer progression.

[12] JA Aronovitz and DR Nelson. Universal features of polymer shapes. Journal de physique,

47(9):1445–1456, 1986.

[20] Kathleen S Metz Reed, Andrew Fritz, Haley Greenyer, Kerstin Heselmeyer-Haddad, Seth Frietze, Janet Stein, Gary Stein, and Tom Misteli. Genome reorganization and its functional impact during breast cancer progression.
